# Expanding Plasma Proteome Coverage While Preserving Biological Signatures: A Systematic Characterization of Enrichment and Preanalytical Variables

**DOI:** 10.64898/2026.11.11.765819

**Authors:** Arthur Viodé, Liliana Malinovska, Sandra Schär, Xaver Wurzenberger, Polina Shichkova, Ina Aretz, Vanessa Buehlmann, Kevin Kraft, Wouter Van Bergen, Nicole Barlow, Leander Runtsch, Christopher R. Below, Yuehan Feng, Amaury Lachaud, Nils A. Kulak, Oliver Rinner, Roland Bruderer

**Author notes:** Correspondence should be addressed to Roland Bruderer, Wagistrasse 21, 8952 Schlieren, Switzerland. These authors contributed equally (Arthur Viodé, Liliana Malinovska, Sandra Schär).

## Abstract

Plasma is a rich and accessible source for biomarkers. The analytical challenge of the steep dynamic range was recently addressed using enrichment technologies. Enrichment technologies have been shown to be sensitive to preanalytical variables, making this an active area of research.

We established the P2 enrichment system as high-performing and robust for large scale studies, with reproducible identifications and precision across batches, days, operators and laboratories. We characterized P2-based enrichment for preanalytical variables (eight blood collection types, centrifugation, freeze-thaw and pathogen inactivation). K_2_EDTA-plasma resulted in the deepest proteome coverage with about 6000 protein groups on the timsTOF HT.

We spiked 16 heavy-labelled proteins into platelet-poor and platelet-contaminated plasma. Importantly, linearity, accuracy, and precision were high in both platelet-poor and platelet-contaminated plasma for both P2-enriched and neat preparations. In a cancer cohort, we demonstrated that 97% of differentially abundant proteins retained the same direction of regulation after P2-based enrichment compared with the neat preparation. Differentially abundant proteins identified in a colorectal cancer pilot cohort remained largely conserved after platelet contamination, including previously reported biomarkers.

We demonstrated that P2-based enrichment enables deep, large scale proteome profiling and preserves biological information despite preanalytical variation. Candidates emerging from P2-based discovery can be validated in independent cohorts with targeted assays.

**Bullet points:**

- We present the P2 based-enrichment workflow and show that it is well suited for deep plasma and serum profiling, standardized, reproducible over labs and time
- The proteome coverage was characterized for eight preanalytical variables of plasma and serum, including centrifugation and pathogen inactivation. Heparin-based plasma and serum gave the lowest proteome coverage
- A P2 Core Proteome of 2662 proteins, which are consistently observed, was established and proposed for usage to QC the enrichment performance
- Linearity of quantification of the P2-based enrichment remains comparable with and without cellular contamination for controlled spike-in experiments and similar effects are observed in neat
- Over 90% of differentially abundant proteins show directionally conservation across neat and P2 based enrichment in a plasma- and serum-based pancreatic cancer study
- Cellular contamination of plasma increases the proteome depth upon enrichment and retains biological quantitative information

## Introduction

Proteins are the main functional molecules of biological systems^1,2^ and alterations in protein abundance, localization, or interactions have been frequently associated with disease onset, progression, and response to therapy^3^. Blood collection is minimally invasive, longitudinal sampling is feasible, and large clinical biobanks are available for retrospective studies. Blood contains an exceptionally rich proteome comprising proteins secreted from tissues, immune cells, extracellular vesicles, and circulating cells^4–6^. Current estimates suggest that several thousand proteins can be detected in plasma, while potentially most proteins may be represented at low concentration levels^7^. Still, comprehensive characterization of the plasma proteome remains challenging, as it spans an estimated dynamic range of more than twelve orders of magnitude. About twenty highly abundant protein families, including albumin, immunoglobulins, transferrin, and haptoglobin, account for more than 99% of the total protein mass^7–9^. In contrast, many proteins with the highest biomarker potential, such as tissue leakage proteins, cytokines, growth factors, and signaling molecules, are present at concentrations several orders of magnitude lower.

Mass spectrometry (MS)-based proteomics has become the leading platform for unbiased and quantitative characterization of complex proteomes^10–12^. Technological advances in sample preparation, liquid chromatography, mass analyzers, and particularly data-independent acquisition (DIA) have substantially improved sensitivity, reproducibility, and throughput^13^. This has enabled the reliable quantification of hundreds to about one thousand proteins from neat plasma and is now applied to large-scale clinical studies^14,15^. Nevertheless, the dynamic range of plasma continues to restrict access to many low-abundance proteins that are considered highly relevant for biomarker discovery, e.g. stemming from the brain^9,16^.

To address this limitation, numerous strategies have been developed to reduce plasma dynamic range prior to MS analysis^17^. Enrichment strategies aim to selectively enrich lower abundance proteins through mechanisms like peptide ligand libraries, protein corona formation, extracellular vesicle enrichment, precipitation or other affinity-based approaches^17–19^. Recent nano and micro particle-based enrichment technologies, including ProteoMiner^20^, Proteograph technology^21^, Mag-Net^22^, ENRICH-iST^17^, P2^17,23^, Proteonano^24^, and related methods, have reported substantial increases in proteome depth, enabling identification of several thousand proteins from plasma^17,19,25–27^. These gains are driven by the protein corona effect^28–31^, which compresses the dynamic range by depleting or reducing abundant proteins and enriching low abundant proteins. Importantly, relative quantification has been shown to be preserved in proteome mixture and single-protein spike-in experiments^18,32,33^.

Increasing evidence suggests that particle-based enrichment technologies are influenced by preanalytical variables^33–38^. Sample collection, anticoagulant choice, processing delays and sample aging, centrifugation protocols, storage conditions, freeze-thaw cycles, and residual cellular contamination from platelets, red blood cells, and peripheral blood mononuclear cells can affect the resulting proteome. Recent studies demonstrated that platelet and blood-cell contamination can increase protein identification by several thousand proteins in enrichment-based workflows and may lead to the misinterpretation of contamination markers as disease biomarkers^33,34,36,37^. To our knowledge, previous contamination studies have largely focused on proteome depth, contamination markers and false biomarker discovery, whereas the influence on preservation of disease-associated fold-change direction has not been systematically evaluated.

P2 is an enrichment technology designed to extend proteomic coverage across plasma and other biofluids while maintaining quantitative robustness and scalability. P2-based enrichment has been successfully tested and used for biomarker discovery and compared to other enrichment technologies^23,39–43^. In this study, we demonstrate that P2-based enrichment is a highly reproducible plasma proteomics workflow suitable for large-scale biomarker discovery studies. The workflow enables deep and robust proteome profiling across a range of preanalytical variables, including blood collection tube type, centrifugation protocols, and pathogen inactivation procedures. We assessed the enrichment quality of the workflow with a newly established P2 Core Proteome panel of reproducibly identified proteins. We further show that cellular contamination increases protein identifications, whereas quantitative accuracy, precision and the detection of disease-associated signals remain largely preserved, both in spike-in experiments with known abundance differences and in clinical cohorts. Together, these results establish P2-based enrichment as a sensitive, scalable, and reproducible approach for plasma biomarker discovery while providing important insights into the impact of preanalytical variation and cellular contamination on proteomic measurements.

## Experimental Procedures

### Material

Chloroacetamide, tris(2-carboxyethyl)phosphine, trifluoroacetic acid, formic acid, ammonium formate, acetonitrile, HPLC water, ammonium bicarbonate, and urea were purchased from SIGMA-Aldrich (Munich, Germany), trypsin sequencing grade was purchased from Promega (Madison, WI, United States), Lys-C was purchased from (KPL Bio, Copenhagen, Denmark), P2-iST Plasma kit (PreOmics, Martinsried, Germany). ^15^N-heavy labeled, recombinant proteins (Table S1) were purchased from (Absea Biotechnology, Berlin, Germany).

### Biofluids samples

The human K_2_EDTA-plasma pool consisted of a pooled plasma sample generated from multiple healthy and diseased donors prepared as K_2_EDTA-plasma and were purchased from Proteogenex (Inglewood, CA, United States) labelled here QC pool. A second K_2_EDTA-plasma pool was purchased from Diaserve Laboratories (Iffeldorf, Germany) labeled here DS30. For the different plasma and serum preparations, specimen from the same 4 donors were collected in various collection tube K_2_EDTA, K_3_EDTA, sodium-citrate, acid citrate-dextrose (ACD), citrate phosphate dextrose (CPD), lithium-heparin, sodium-heparin and serum. For K_2_EDTA samples, plasma was additionally prepared using three processing protocols: a single-spin protocol, a double-spin protocol (2x 1,500 × g for 15 min), and the UK Biobank plasma collection protocol. All were purchased from Research Donors (London, United Kingdom). K_2_EDTA platelet rich (PRP) and poor (PPP) plasma were purchased from Research Donor. They consist of a pool of 2 donors. Platelet rich and poor plasma was prepared according to Korff et al^34^. Briefly, whole blood was centrifuged at 500 × g for 7 min to obtain platelet-rich plasma. Half of the PRP was stored at -20 °C or -80 °C. The remaining PRP was centrifuged again at 500 × g for 7 min, and the recovered supernatant was subjected to two sequential centrifugation steps at 3,000 × g for 7 min. The final supernatant was collected as platelet poor plasma and stored at -20 °C or -80 °C. The pellet was washed twice with 4 mL PBS/EDTA by centrifugation at 3,000 × g for 7 min. The resulting pellet was collected as purified platelets. To isolate erythrocytes, the cellular pellet obtained after the initial blood centrifugation step was centrifuged at 3,000 × g for 7 min. The supernatant, and the uppermost layer of the erythrocyte were carefully removed and discarded. The remaining erythrocytes were washed twice with PBS/EDTA by centrifugation at 3,000 × g for 7 min, followed by removal of the supernatant and the upper cell layer after each wash. 24 K_2_EDTA -plasma samples from 12 colorectal cancer and 12 healthy donors were purchased from Proteogenex. For the analysis, they were spun 30 min at 3000 x g at 4 °C and then the supernatant was taken to get platelet-poor plasma. For the platelet spike in experiment, equivalents of estimated 10^6^ and 10^7^ cells were mixed with the spun plasma prior to P2-based enrichment.

### Neat preparation

The equivalent of 1 µL of neat plasma or serum samples were processed using protein aggregation capture (PAC) in a semi-automated fashion using a KingFisher Flex (Thermo Scientific). The PAC protocol was performed as described previously ^44^. In brief, samples were mixed with carboxylate-modified Sera-Mag SpeedBead magnetic particles (Sigma-Aldrich), and protein aggregation was induced by the addition of methanol to achieve a final concentration of 70% (w/w). Beads were washed twice with a 95% acetonitrile solution and once with 70% ethanol. Proteins were reduced and alkylated using reduction/alkylation buffer (0.1 M ammonium bicarbonate (ABC), 20 mM tris(2-carboxyethyl)phosphine, and 80 mM chloroacetamide (CAA), 70% ethanol) for 60 min at 37 °C. Samples were digested with digestion buffer (50 mM tris(hydroxymethyl)aminomethane, trypsin (1:50 enzyme to substrate), and Lys-C (1:200 enzyme to substrate)) for 18 h at 37 °C. After beads removal samples were acidified with 20% trifluoroacetic acid (TFA) and subjected to peptide cleanup.

### P2-based enrichment sample preparation

P2-based enrichment was performed either following the Biognosys contract research method P2-PAC (Biognosys AG, Schlieren, Switzerland) or using the P2-iST Plasma kit (PreOmics GmbH, Martinsried, Germany) following the manufacturer’s instructions. The protein corona formation step was identical, and the overall performance was found to be very similar (Figure S1) and Table S2 details which approach was used for which figure. Briefly for P2, 100 µL of biofluids were incubated with 75 µL of binding buffer (EDTA-plasma binding buffer or serum/heparin optimized binding buffer for serum and heparin, Biognosys AG, Schlieren, Switzerland) and 25 µL of P2 single particle solution for 40 min at 30°C. Samples were processed in a semi-automated fashion using a KingFisher Flex (Thermo Scientific, Waltham, MA, United States). Digestion was performed for 18 h at 37 °C using 1 µg trypsin (Promega) and 0.5 µg of Lys-C (KPL Bio). Digestion was stopped by the addition of 10 µL of 20% TFA and samples were subjected to peptide cleanup.

### Peptide cleanup

Peptide cleanup and desalting were performed using an Oasis hydrophilic–lipophilic balanced (HLB) µElution Plate (Waters). HLB plates were conditioned by the sequential addition of 100% acetonitrile (ACN) and washing buffer (1% ACN, 0.1% TFA in water). Peptides were loaded onto the conditioned HLB plates and washed twice with the washing buffer. Peptides were eluted using an elution buffer (30% ACN, 0.1% TFA in water) and dried down using a SpeedVac (Thermo Scientific). Peptides resulting from P2-iST Plasma solution were cleaned up following the manufacturer’s instructions. Dried peptides were resolubilized in resuspension buffer (1% ACN, 0.1% FA in water with iRT peptides).

### Cellular contamination spiking into plasma

Platelets and erythrocytes were obtained as described above. The cell count was estimated from the donor counts before collection (platelets 197x10^9^/L; erythrocytes 4.61x10^12^/L) with 40% and 25% loss, respectively. For the controlled spike-in contamination model, equivalents of 10^5^, 10^6^, 10^7^ and 10^8^ cells were added to platelet-poor plasma. For platelet-rich plasma supplementation, fractions of 5%, 10%, 25%, 50% and 75% were added to platelet-poor plasma. The samples were processed as neat and P2-based enrichment preparations as described above.

### Heavy labelled protein spiking into plasma

Heavy isotope proteins were spiked into the K_2_EDTA-plasma QC pools at defined concentrations of a linear dilution, close to their endogenous levels in triplicates (Table 1). Next, the mixtures were prepared both with the neat workflow and with P2-based enrichment, as described above. One non-spike sample was used as blank measurement.

**Table 1.** Recombinant heavy labelled protein spike-in. Endogenous levels and concentration layout

| Protein | endogenous concentration pg/L | Spike-in concentration [ng/µL] |  |  |  |  |  |
|---|---|---|---|---|---|---|---|
|  |  | C1 | C2 | C3 | C4 | C5 | C6 |
| CALM1 | 6.70E+07 | 2.080 | 1.040 | 0.520 | 0.260 | 0.130 | 0.070 |
| CTSD | 1.40E+08 | 4.520 | 2.260 | 1.130 | 0.570 | 0.290 | 0.150 |
| CTSS | 1.40E+07 | 2.060 | 1.030 | 0.520 | 0.260 | 0.130 | 0.070 |
| GSR | 3.20E+07 | 2.080 | 1.040 | 0.520 | 0.260 | 0.130 | 0.070 |
| HBD | 8.70E+09 | 18.470 | 9.230 | 4.620 | 2.310 | 1.160 | 0.580 |
| IGFBP4 | 2.60E+08 | 5.590 | 2.790 | 1.400 | 0.700 | 0.350 | 0.180 |
| MGP | 2.00E+07 | 2.330 | 1.170 | 0.590 | 0.300 | 0.150 | 0.080 |
| NAP1L1 | 1.30E+07 | 2.010 | 1.010 | 0.510 | 0.260 | 0.130 | 0.070 |
| PPBP | 3.80E+09 | 13.930 | 6.960 | 3.480 | 1.740 | 0.870 | 0.440 |
| SAR1A | 8.10E+05 | 0.780 | 0.390 | 0.200 | 0.100 | 0.050 | 0.030 |
| TUBA8 | 1.10E+08 | 4.170 | 2.080 | 1.040 | 0.520 | 0.260 | 0.130 |
| TPM1 | 1.30E+08 | 2.080 | 1.040 | 0.520 | 0.260 | 0.130 | 0.070 |
| APOE | 1.70E+10 | 19.990 | 9.990 | 5.030 | 2.510 | 1.140 | 0.000 |
| S100A9 | 1.10E+09 | 12.510 | 6.250 | 3.140 | 1.570 | 0.710 | 0.000 |
| ENO2 | 4.70E+07 | 2.510 | 1.250 | 0.630 | 0.320 | 0.160 | 0.070 |
| FTH1 | 2.80E+06 | 1.250 | 0.620 | 0.310 | 0.160 | 0.080 | 0.040 |

### LC-MS acquisition

800 ng of peptides per sample were loaded onto a Evotip Pure (Evosep, Cat.-No.: EV2011) following the manufacturer’s instructions. Peptides were separated on an IonOpticks Aurora series Elite CSI 75 µm x 150 mm C18 reversed phase column (AUR4-15075C18-CSI) using an Evosep One liquid chromatography system operated with the Whisper-Zoom 40 SPD method. The LC was connected to a Bruker Daltonics timsTOF HT mass spectrometer equipped with a CaptiveSpray 2 ion source. Data were acquired using diaPASEF. The method consisted of one full range MS1 scan from 100 –1700 m/z with an applied ion mobility range from 0.77 –1.18 1/k0 with ramp and accumulation times set to 70 ms (100 % duty cycle) and 13 PASEF ramps. During the operation of the instrument, the mobility was calibrated every 12h to ensure proper acquisition of the data.

### Mass Spectrometric Data Analysis

Raw data were processed using Spectronaut^45^ in directDIA mode. The canonical human reference proteome UP000005640 from 2026-01-01 was used. The search allowed for 2 missed cleavages and variable modifications (N-term acetylation, ammonia-loss, and oxidation (MP). For identification, the false discovery rate on peptide and protein level was set to 1%. Protein quantities were calculated using the maxLFQ approach^46^. Differential abundance testing was set to two sample t-test, not assuming equal variance, two sided and using all quantitative information levels^47^. Multiple testing correction for false discovery rate was done using the Storey approach^48^.

For recombinant protein spike-in experiments, analysis in Spectronaut was performed with a labelled workflow using the labels 15N(1), 15N(2), 15N(3) and 15N(4) and the settings described above.

### Experimental Design and Statistical Rationale

All data analyses were conducted in the R programming language (V. 4.5.1) using the tidyverse (V. 2.0.0) package environment for data handling and modification in RStudio (V. 2025.05.1+513) as well as Python 3.10.11. Prior to data handling, the quantitative data was extracted on peptide precursor level from Spectronaut using a custom report schema. The differentially abundant proteins were filtered with an FDR <1%, a log_2_ fold change of >0.58 and proteins with >2 stripped sequences. The coefficient of variation plots are in percent and cut at 100%, for better visibility of the differences. The number of omitted values is indicated.

Assessment of different protein group identifications and coefficients of variation was modelled as linear mixed-effects models using the following formula (Table 2).

**Table 2:**
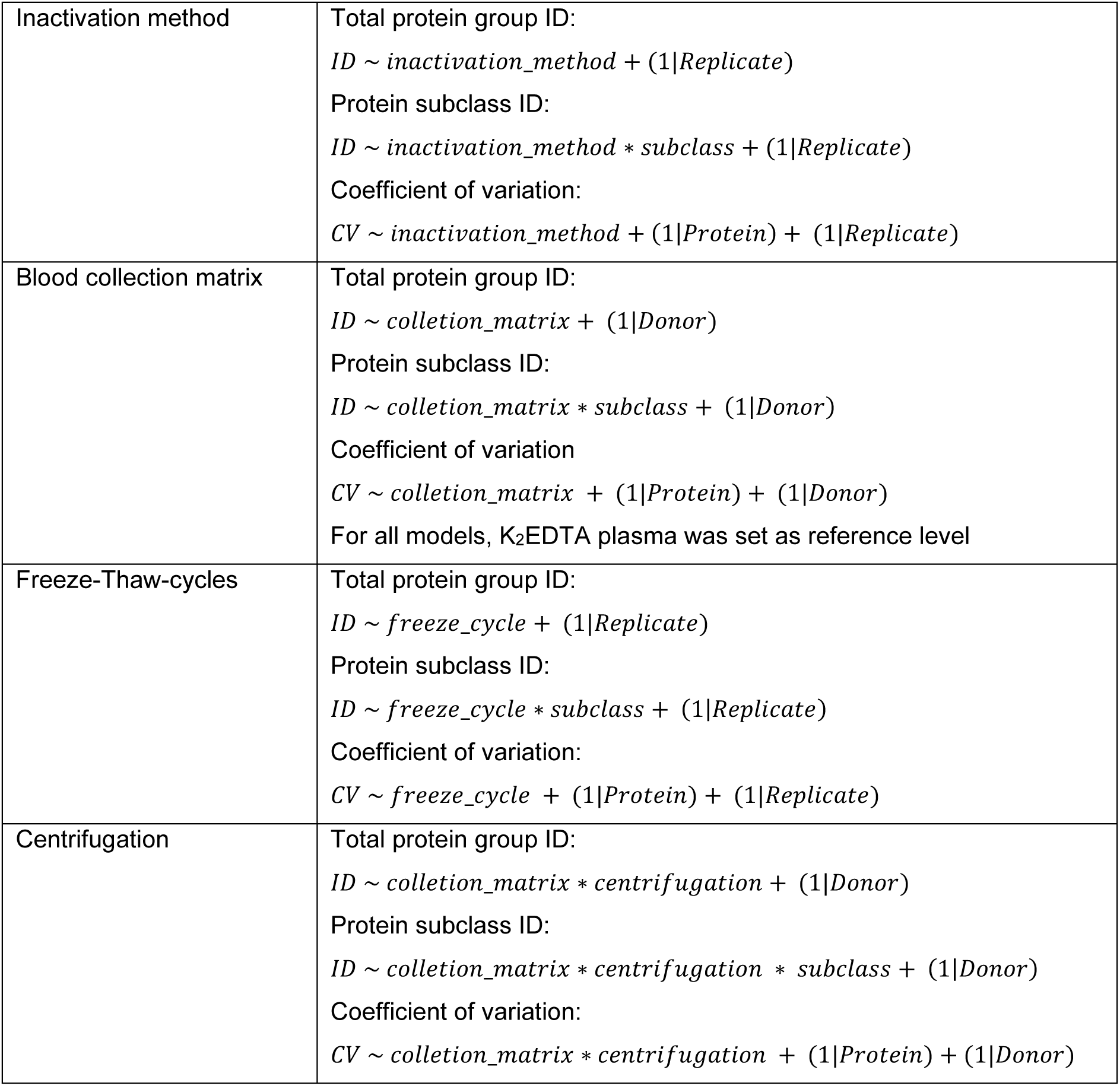
Mixed effect models for preanalytical variables. Assessment of protein group identifications and coefficients of variation were modelled as linear mixed-effects models

|  |  |
|---|---|
| Inactivation method | Total protein group ID:<br>ID ∼ inactivation_method + (1\|Replicate)<br>Protein subclass ID:<br>ID ∼ inactivation_method ∗ subclass + (1\|Replicate)<br>Coefficient of variation:<br>CV ∼ inactivation_method + (1\|Protein) + (1\|Replicate) |
| Blood collection matrix | Total protein group ID:<br>ID ∼ colletion_matrix + (1\|Donor)<br>Protein subclass ID:<br>ID ∼ colletion_matrix ∗ subclass + (1\|Donor)<br>Coefficient of variation<br>CV ∼ colletion_matrix + (1\|Protein) + (1\|Donor)<br>For all models, K<sub>2</sub>EDTA plasma was set as reference level |
| Freeze-Thaw-cycles | Total protein group ID:<br>ID ∼ freeze_cycle + (1\|Replicate)<br>Protein subclass ID:<br>ID ∼ freeze_cycle ∗ subclass + (1\|Replicate)<br>Coefficient of variation:<br>CV ∼ freeze_cycle + (1\|Protein) + (1\|Replicate) |
| Centrifugation | Total protein group ID:<br>ID ∼ colletion_matrix ∗ centrifugation + (1\|Donor)<br>Protein subclass ID:<br>ID ∼ colletion_matrix ∗ centrifugation ∗ subclass + (1\|Donor)<br>Coefficient of variation:<br>CV ∼ colletion_matrix ∗ centrifugation + (1\|Protein) + (1\|Donor) |

For the recombinant protein spike in experiment, limit of detection (LOD) and limit of quantification (LOQ) for each peptide was estimated as described previously^49^. Heavy protein quantities were calculated as mean intensities of the top 3 heavy peptide intensities per condition, after excluding peptides below the estimated LOQ. Linearity was assessed for each protein in each contamination background using a linear regression model (i) and the adjusted coefficient of determination (adj. R^2^) was obtained.

(i) log_2_(*Protein quantity*) ∼ log_2_(*Concentration*)

A threshold of adj. R^2^ > 0.8 was set for a good linear response. Slopes were estimated using a linear mixed-effect model (ii) and slope differences were assessed as ANOVA of the interaction terms.

(ii) log_2_(*Protein quantity*) ∼ log_2_(*Concentration*) ∗ *Conamination*_*background* + (1|*Replicate*)

Endogenous peptide levels were estimated from the light to heavy ratios of peptides above LOQ as described previously^50^ as weighted linear regression (iii)

(iii) log_2_(*Concentration*) ∼ 0 + *Ratio*

Endogenous protein levels were calculated as mean endogenous peptide levels, differences in protein levels across contamination backgrounds were assessed using ANOVA.

## Results

### Robust clinical applicability of the P2-based enrichment with sensitivity to pre-analytical sample treatment

An important prerequisite for large-scale clinical studies is that plasma proteomics measurements remain reproducible over time and across analytical batches. We therefore investigated whether the P2-based enrichment maintains consistent analytical depth and quantitative performance over 1.5 years routine operation across different P2 particle batches, operators, days, and laboratories.

To evaluate the robustness of the P2-based enrichment, we prepared a K_2_EDTA-plasma QC pool that was used throughout the study. We first assessed P2 batch variability by processing the plasma pool with four independent P2 particle batches. Protein identifications were highly consistent across batches with 6352 ± 32 protein groups while median protein group CVs ranged from 8.0% to 9.0% within individual batches (Fig. 1A). Across all batches, the median CV was 10.8%. Inter-day reproducibility was also maintained, with 5859 ± 24 protein groups identified and a median protein group CV of 15.2% (Fig. 1B). Inter-operator reproducibility was similarly high, with 6055 ± 7 protein group identifications and a median protein group CV of 10.8% across operators (Fig. 1C).

**Figure 1.**
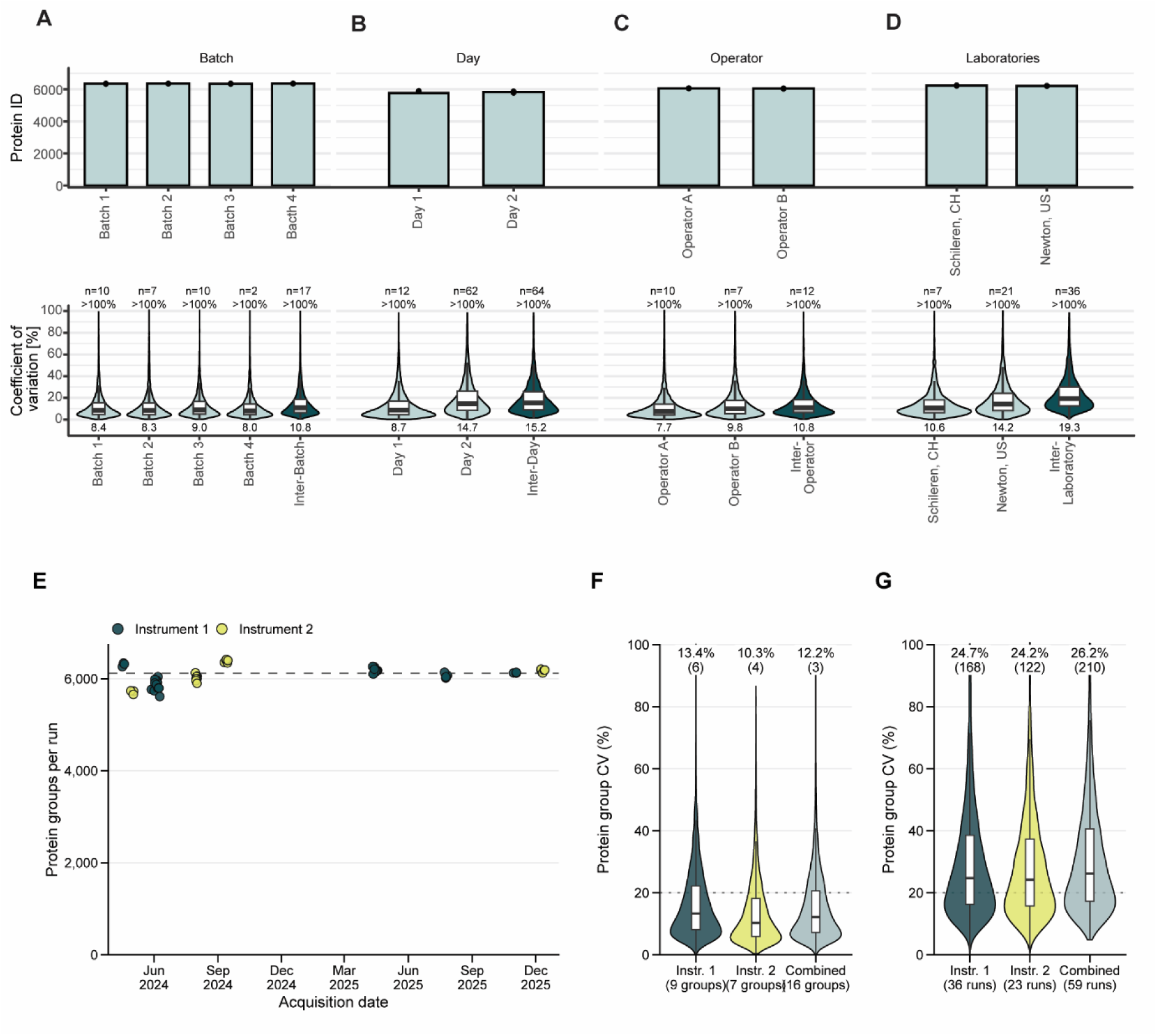
Reproducibility and long-term performance of the P2 plasma workflow. (A-D) Comparison of Protein Group Identifications (upper plot) and coefficient of variation (CVs; lower plot) using K_2_EDTA-plasma QC pool processed (A) with four different P2 particle batches (n=4), (B) by two operators (n=3), (C) on two different days (n=3); and (D) at two laboratories (n=3). Mean protein group identifications are displayed as bars, with individual measurements shown as dots. CVs were calculated on protein group level for intra-sample variability (light teal) and inter-sample variability (dark teal) and are displayed as violin plots with overlaid boxplot. Mean CV values are indicated at the bottom. Outliers (>1.5 x interquartile range) are not displayed. CVs >100% are excluded, the number of excluded observations is indicated at the top. **(E)** Scatterplot displaying protein group identifications from 59 measurements of the K_2_EDTA QC plasma pool across 13 sequences acquired between April 2024 and December 2025 on two timsTOF HT instruments (dark teal, instrument 1, 36 runs; yellow, instrument 2, 23 runs). The dashed line indicates the median number of protein group identifications (6123 protein groups). (**F**) Within-sequence CVs calculated across replicate measurements of K_2_EDTA QC plasma pool within individual P2 enrichment experiments, with 16 replicate groups and 3–8 measurements per group. (**G**) Long-term CVs calculated across all measurements acquired between 2024 and 2025 on two timsTOF instruments, including independent P2 preparations, two LC gradients and 20 months of operation.

To assess inter-laboratory reproducibility, the P2-based enrichment was independently executed in Biognosys laboratories in Schlieren, Switzerland, and Newton, USA. The two laboratories identified 6209 and 6235 protein groups, respectively, with a CV of 10.6% and 14.2%, respectively (Fig. 1D). Comparison across laboratories resulted in a median protein group CV of 19.3%, demonstrating robust and reproducible protein quantification across sites.

To further assess long-term robustness under routine laboratory conditions, we performed a combined analysis of P2 preparations generated over a 1.5-year period using five P2 particle batches, three operators, and two LC-MS instruments. All analyses were performed using aliquots of the same pooled K_2_EDTA QC plasma pool. Across all preparations, a median of 6123 ± 202 protein groups was identified, of which 4762 were consistently detected (Fig.1E) with a median CV of 12.2% for samples within acquisition sequences of one individual project (Fig. 1F) and 26% across all acquisitions (Fig. 1G).

Combined, these results demonstrate high repeatability of protein identification and robust quantitative precision, maintaining median protein group CVs below 20% across individual assessments of batch, operator, day and laboratory variability and at 26% when all these variables were combined over two years of routine operation. These findings demonstrate that P2 provides highly stable and reproducible plasma proteome profiling over time, supporting its application in large-scale biomarker discovery and clinical studies.

For application in large-scale clinical studies, the P2-based enrichment workflow must also be compatible with sample treatments that are routinely applied to plasma specimens prior to proteomic analysis. In studies involving infectious diseases, plasma specimens may contain pathogenic viruses, bacteria, or parasites and therefore often require pathogen inactivation prior to downstream analysis. We therefore investigated whether commonly used inactivation procedures affect the analytical depth, quantitative precision, and overall performance of the P2-based enrichment workflow. For inactivation of bacteria, viruses and parasites, we assessed filtration using 0.22 µm filters^51^ as well as heat-inactivation at 56 °C for 1 hour^52^. For inactivation of enveloped viruses, we tested incubation with a surfactant (1% and 2% Triton-X100)^53^. We observed a reduction in protein group identifications for all inactivation methods applied to the K_2_EDTA QC plasma pool (Fig. S2A), with filtration and heat-inactivation yielding the lowest identifications (about 40% lower). Interestingly, the effect of inactivation was not uniform across protein groups particularly relevant to plasma proteomics, such as growth factors, transcription factors and cytokines. For Triton X-100 treatment, identification of cytokines, growth factors, and transcription factors were largely unaffected, therefore it is a well-suited inactivation if applicable. (Fig. S2B). In addition, we observed an increase in variability for all inactivation methods. Heat inactivation had the greatest impact on those protein classes (Fig. S2C). Together, these findings demonstrate that the tested pathogen inactivation procedures reduce proteome coverage and moderately impair quantitative precision in the P2-based enrichment workflow. This highlights the need to carefully consider sample inactivation strategies in plasma proteomics studies.

### Pre-analytical sample processing influences P2-based enrichment performance

Enrichment technologies for plasma and serum proteomics are often optimized and benchmarked on rather uniform datasets and selected preparation protocols. Their application, however, is aimed at large scale studies, where sample collection can be subject to major variabilities. Sample collection method, anticoagulant choice, processing delays, centrifugation protocols, storage conditions, freeze-thaw cycles represent pre-analytical variables which can impact protein identification performance.

We systematically assessed the impact of blood collection methods such as plasma with the anticoagulants K_2_EDTA, K_3_EDTA, sodium-citrate, acid-citrate-dextrose (ACD), citrate-phosphate dextrose (CPD), lithium-heparin, or sodium-heparin. Importantly, blood from four donors was collected in parallel into all the eight collection types (Fig. 2A). We observed significant variations in protein group identification (Fig. 2B). The blood collection method had a highly significant effect on protein group identifications (p < 0.0001). Compared to K_2_EDTA, which was used as reference matrix, all other collection methods yielded significantly fewer protein group identifications, with the strongest reductions observed for lithium-heparin (−42%) and sodium-heparin (−47%), while CPD (−7.1%) and K_3_EDTA (−7.0%) showed the smallest performance differences. Next, we assessed the effect of the blood collection method on the identification of protein groups particularly relevant to plasma proteomics using a linear mixed-effects model and concluded that the effect of collection method is not uniform across the protein classes (p = 3.95 × 10⁻^8^). Transcription factors appear particularly sensitive to the collection method compared to growth factors and cytokines that were mostly reduced in heparin-based plasma (Fig. 2C). Overall, we conclude that the highest proteome coverage is achieved using K_2_EDTA plasma and the use of other blood collection methods will influence the coverage to different extents. Interestingly, this effect of the collection method is more pronounced in the enrichment technology, as neat plasma analyses from the same samples do not show a significant effect on protein groups identification (p = 0.31) (Fig. S3A). Importantly, P2-based enrichment still yielded substantially higher protein identifications across all tested collection methods.

**Figure 2:**
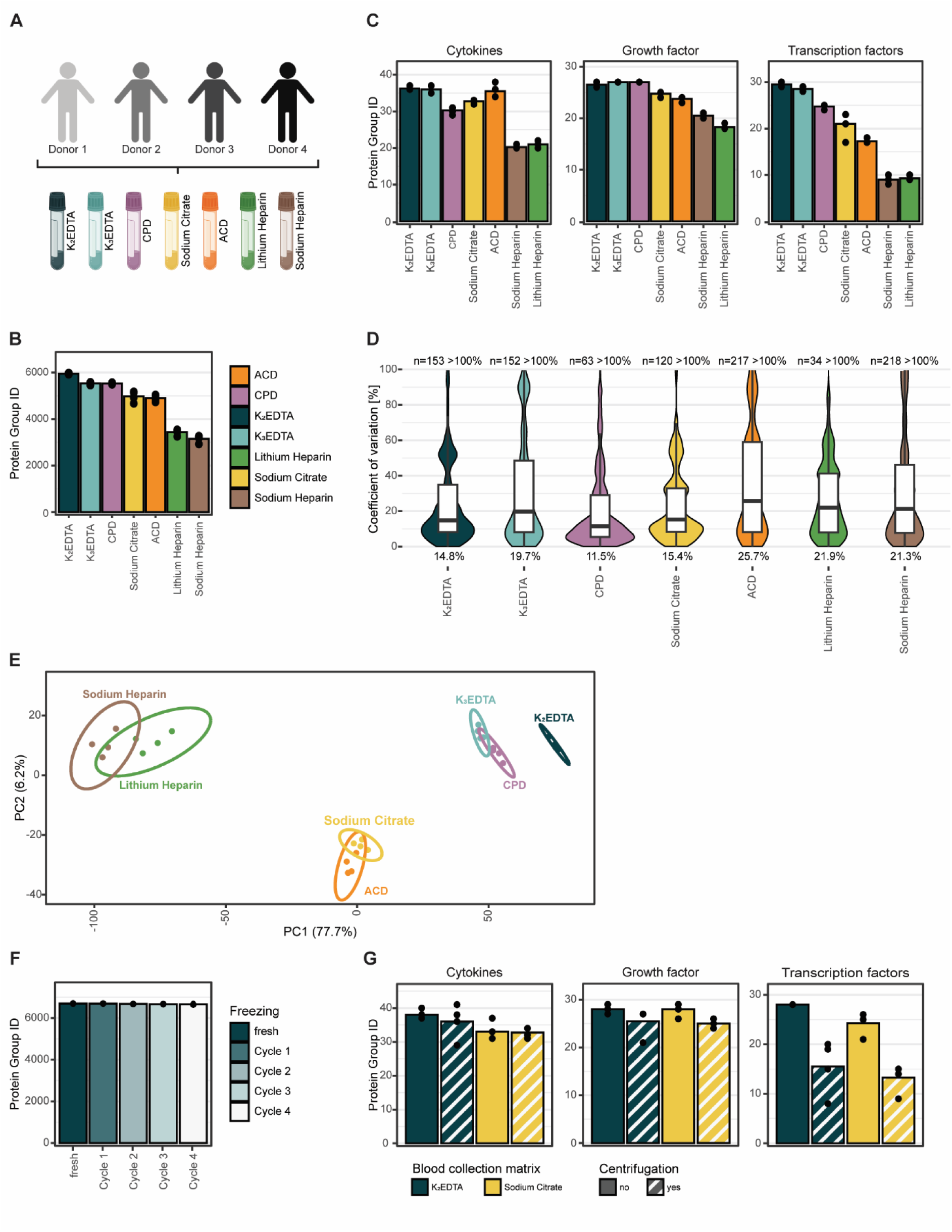
Effect of pre-analytical variables on P2-based enrichment performance. **(A)** Schematic representation of the blood collection strategy. Blood was collected from 4 different donors, using each of the seven blood collection matrices (dark teal, K_2_EDTA; light teal, K_3_EDTA; purple, CPD; yellow, sodium-citrate; orange, ACD; green, lithium-heparin; brown, sodium-heparin). Image generated using BioRender.com (biorender.com). (**B**) Comparison of protein group identifications following P2-based enrichment of plasma collected as described in (A). Mean protein group identifications are displayed as bars, with individual measurements shown as dots. **(C)** Protein group identification for selected protein classes (cytokines, growth factors, transcription factors) in the samples described in (A). **(D)** Coefficients of variation (CVs) calculated at the protein group level for the samples described in (A) and displayed as violin plots with overlaid boxplots. Mean CV values are indicated at the bottom. Outliers (>1.5 x interquartile range) are not displayed. CVs >100% are excluded; the number of excluded observations is indicated at the top. **(E)** Principal component analysis (PCA) of the protein group quantities in the samples described in (A). **(F)** Comparison of protein group identifications in K_2_EDTA-plasma subjected to zero to four consecutive freeze thaw cycles (dark teal to white). Mean protein group identifications are displayed as bars, with individual measurements shown as dots. **(G)** Comparison of protein group identifications in K_2_EDTA (dark teal) and sodium-citrate (yellow) plasma from four donors before and after an additional centrifugation step at 2000 x g (striped bar). Protein group identifications are shown for selected protein classes (cytokines, growth factors, transcription factors), mean protein group identifications are displayed as bars, with individual measurements shown as dots.

Next, we assessed if the choice of blood collection method affects the quantitative performance of technology. For biological replicates, we compared the CV of the same set of proteins across the collection types and four donors (Fig. 2D). We observed that the blood collection method significantly affected protein-level quantitative precision (p < 0.0001). K_2_EDTA, CPD, and K_3_EDTA exhibited the lowest coefficients of variation, whereas ACD-, citrate-, and particularly heparin-based matrices showed increased variability. To investigate the variance across the collection types, a principal component analysis (PCA) demonstrated that blood collection type introduces quantitative differences, but these differences were highly reproducible within each collection type (Fig. 2E, and S3B). In addition to plasma, we also tested the performance of P2 in serum samples and compared protein identification (Fig. S3C). We observed a consistent decrease in protein identifications, both for total protein groups and for specific protein subclasses, albeit the decrease in numbers being dependent on the subgroup (Fig. S3D). Taken together, we conclude that the choice of blood collection method will have a substantial effect on both analytical depth and quantitative performance when using P2.

Other pre-analytical variables that can be controlled and considered during study design and assessed systematically are freeze-thaw cycles and centrifugation protocols. With a K_2_EDTA-plasma QC pool, we tested up to four freeze-thaw cycles. We observed very little variation in protein group identifications (∼0.6%, p = 0.014) (Fig. 2F). We observed a significant effect (p < 0.0001) of freezing on protein quantification precision (Fig. S4A). After three freeze-thaw cycles, the CV increased by ∼25%, which suggests avoiding more than two freeze-thaw cycles (∼10% CV increase). Next, we evaluated the effects of additional centrifugation cycles to the standard blood collection strategy in K_2_EDTA and sodium-citrate-plasma pools and their influence on analytical proteome depth and quantitative performance. We observed a decrease in protein group identifications (K_2_EDTA: 26.5 %, sodium-citrate: 29.1 %) as a result of centrifugation mediated removal of cells and debris (Fig. S4B). Interestingly, the extent of the loss was not uniform across protein classes (Fig. 2G). For protein group identification, transcription factors showed a highly significant reduction averaged across collection methods (post-hoc contrast, p < 0.0001, ∼45%), while growth factors (post-hoc contrast, p = 0.526, ∼10%) and cytokines (post-hoc contrast, p = 0.133, 1-5%) remain largely unchanged. Centrifugation also significantly affects quantitative precision on protein level (p < 0.0001) for both plasma types (Fig. S4C).

Overall, we confirm that pre-analytical variables influence the performance of the P2-based enrichment technology, and the extent of the effect varies between protein classes relevant to plasma proteomics such as cytokines, transcription factors or growth factors.

### Establishment of a P2 Core Proteome and application as a practical, robust quality metric for enrichment performance

We confirmed that pre-analytical variables can influence P2-based enrichment outcomes. These factors are often difficult to control or assess independently due to the limited availability of matched samples. Therefore, we sought to establish an orthogonal measure of enrichment performance that is applicable to large-scale and longitudinal studies. Our evaluation focused on analytical depth and quantitative precision as more robust metrics of enrichment performance. For this, we established a reference panel of proteins consistently measurable in EDTA plasma, the most used plasma specimen type.

By comparing identified proteins across various in-house studies, we defined a P2 Core Proteome comprising 2662 proteins that were consistently (>50%) identified irrespective of EDTA-plasma source (Table S3). Comparison with the Human Plasma Protein Atlas (V21 14^th^ Oct. 2026) revealed an overlap of 1788 proteins (Fig. 3A). These proteins covered more than eight orders of magnitude in reported plasma concentration, ranging from 4.2 ng/L to 440 mg/L. This demonstrates that the P2 Core Proteome is not limited to the highest-abundance plasma proteins but includes proteins across a broad abundance spectrum.

**Figure 3:**
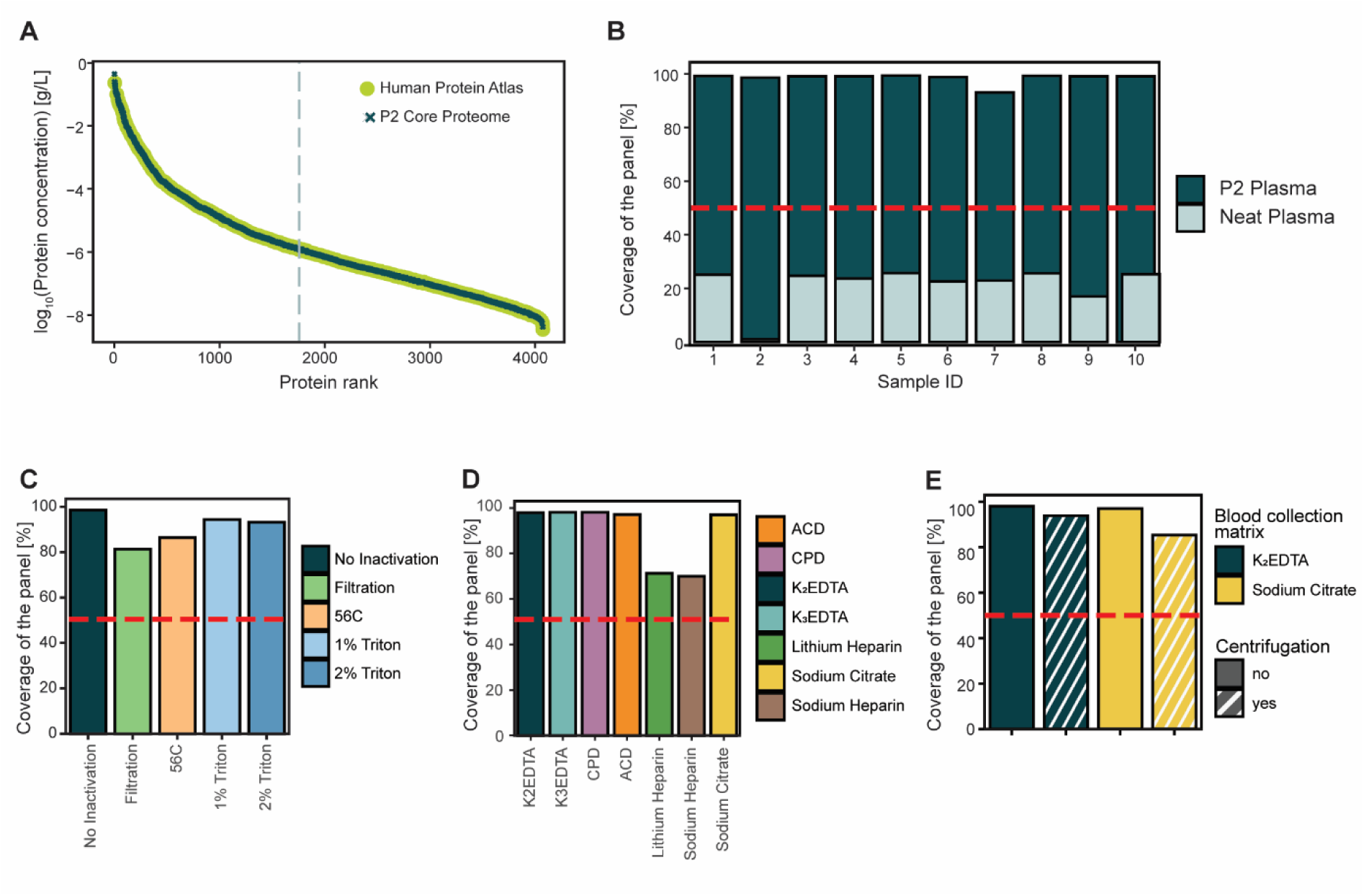
P2 Core Proteome and its application to evaluate the quality of enrichment. (**A**) P2 Core Proteome proteins mapped to protein concentrations reported in the Human Proteome Atlas (HPA) v21^7^and ranked by concentration. The dashed vertical line indicates the median rank of P2 Core Proteome proteins in the Human Proteome Atlas (median = 1756.5). **(B)** Coverage of the P2 Core Proteome in P2-enriched plasma (dark teal) and neat plasma (light teal) across 10 independent samples from the pancreatic cancer cohort. **(C–E)** Coverage of the P2 Core Proteome across different pre-analytical conditions. (C) P2 enrichment of a K₂EDTA plasma QC pool without inactivation pre-treatment (dark teal) or following inactivation by filtration through 0.22 µm filters (green), heat treatment at 56 °C for 1 h (orange), or treatment with 1% or 2% Triton X-100 (light blue and dark blue, respectively). (D) P2 enrichment of plasma collected from four donors using different blood collection matrices (dark teal, K_2_EDTA; light teal, K_3_EDTA; purple, CPD; yellow, sodium-citrate; orange, ACD; green, lithium-heparin; brown, sodium-heparin). (E) P2 enrichment of K_2_EDTA (dark teal) and sodium-citrate (yellow) plasma from four donors before and after an additional centrifugation step at 2,000 × g (striped bars). In (B–E), the red dashed line indicates the predefined quality threshold of 50% P2 Core Proteome coverage.

To investigate the biological characteristics of the P2 Core Proteome, we performed Gene Ontology overrepresentation analyses using the complete set of proteins identified across all P2-based enrichment studies as the background (8839 proteins). This analysis revealed an overrepresentation of cellular components and molecular functions associated with vesicles, microparticles and cell adhesion, as well as biological processes related to wounding, immune effectors process and platelet activation (Fig. S5A). These findings indicate that proteins involved in extracellular vesicle biology and hemostatic and immune-related functions are consistently detected in EDTA-plasma using the P2-based enrichment. Given the overrepresentation of vesicle-associated proteins, we next compared the P2 Core Proteome with the ExoCarta Top-100 proteins (v6, downloaded 14 Oct 2026)^54^. We found that 93 of the 100 ExoCarta proteins were present in the P2 Core Proteome, of which 67 were also included in the Human Plasma Protein Atlas (Fig. S5B). Together with the Gene Ontology analysis, the results indicate that the P2-based enrichment consistently enriches proteins associated with extracellular vesicles including exosomes (30-150nm) and ectosomes/micro vesicles (100nm - 1um)^23^.

Next, we evaluated the utility of the P2 Core Proteome as a quality metric for P2-based enrichment performance. To this end, we assessed the proportion of P2 Core Proteome proteins detected in an independent dataset comprising ten plasma specimens of a pancreatic cancer cohort. Across the samples processed with the P2-based enrichment workflow, the coverage was highly consistent, ranging from 93% to 99% (Fig. 3B), while the identified protein groups exhibited a higher variability, ranging from 4199 to 6089 protein groups (Fig. S5C). In contrast, analysis of neat plasma from the same donors yielded substantially lower and more variable coverage, ranging from 1% to 25.7% (Fig. 3B) and following the fluctuations of the identified protein groups (Fig. S5D). Based on these observations, we defined a P2 Core Proteome coverage threshold of 50% as a quality control criterion for the performance of enrichment. This threshold is approximately two-fold higher than the highest coverage observed in neat plasma and thus effectively discriminates successful P2-based enrichments from non-enriched plasma samples.

With this quality control metric, we evaluated enrichment performance across the different plasma collection, processing, and sample preparation conditions investigated in this study. The different inactivation methods also yielded lower coverage, which was reduced from 98.5% to 81.4% for filtration, 86.4% for heat inactivation and 94.4% and 93.2% for incubation with a surfactant (1% and 2% Triton-X, respectively) (Fig. 3C). Across the blood collection methods, we observed high coverage ranging from 93% to 98% in most preparations except for sodium-heparin (70%) and lithium-heparin (71.2%) (Fig. 3D). Centrifugation of K_2_EDTA- and sodium-citrate-plasma led to a small decrease in coverage from 97.97 % and 96.99 %, respectively, to 93.8% and 85.3% (Fig. 3E). Importantly, all tested conditions remained above the predefined quality threshold of 50% P2 Core Proteome protein coverage. Thus, although several pre-analytical variables decreased the total number of identified proteins (Fig. 2 and Fig. S2), they did not compromise the overall quality of P2-based enrichment.

To assess the generalizability of the enrichment performance score, we applied the P2 Core Proteome coverage metric to the independent plasma proteomics dataset generated by Völlmy et al. using different workflows and instrumentation^23^. Despite methodological and analytical differences, P2 Core Proteome coverage remained consistently high, ranging from 95.6 to 99.5% across most sample preparation conditions (Fig. S6). The only exception was ultracentrifuged plasma, where the coverage decreased substantially to 52.5%-58.7%, yet remained above the predefined threshold. As ultracentrifugation is designed to deplete extracellular vesicles from plasma^55^, this reduction is consistent with the observed enrichment of extracellular vesicle-associated proteins within the P2 Core Proteome. These findings demonstrate that the P2 Core Proteome coverage metric is transferable across studies and analytical platforms.

Taken together, these findings establish P2 Core Proteome coverage as a robust and biologically informed metric for assessing P2-based enrichment performance. The metric remained stable across diverse pre-analytical conditions, analytical workflows, and mass spectrometry platforms, while sensitively detecting conditions known to affect the extracellular vesicle-associated protein fraction. As such, P2 Core Proteome coverage provides a practical quality control measure for standardized application of the P2-based enrichment in large-scale and longitudinal plasma proteomics studies.

### A controlled spike-in model enables assessment of contamination effects

The observed loss of protein identifications upon centrifugation has been reported previously and have been attributed to removal of platelet-, erythrocyte-, and coagulation-derived cellular material^36,37^. Controlled addition of contaminants such as platelets or erythrocytes has been shown to affect proteome depth and quantitative measurements, particularly in particle-based enrichment^36,37^. The presence of such contaminants in plasma samples has therefore been suggested to affect quantification accuracy and precision and confound biomarker discovery in plasma proteomics. To investigate this, we established a controlled spike-in model, in which defined levels of different contaminating materials (platelets, red blood cells or platelet-rich plasma) were added to platelet-poor K_2_EDTA-plasma, generating samples that span a range of contamination levels. Consistent with previous reports, increasing contamination resulted in a progressive increase in protein group identifications (Fig. S7A). We further quantified contamination using the platelet and erythrocyte contamination indices introduced by Korff et al.^34^ (Fig. S7B). Matching these samples against the P2 Core Proteome revealed distinct effects of different contamination sources on the enrichment quality scores (Fig. S7C). Erythrocyte contamination yielded enrichment quality scores ranging from 78.2% to 87.7%, with score fluctuations closely mirroring changes in protein group identifications. In contrast, supplementation with platelet-rich plasma increased the enrichment quality score from 81.8% in platelet-poor plasma to 98.7% at 5% platelet-rich plasma. Further increases in platelet-rich plasma content had little additional effect on enrichment quality, with scores remaining between 98.3% and 99.1%, despite continued increases in protein group identifications. Platelet contamination, however, resulted in concurrent increases in both enrichment quality score and protein group identifications. These findings suggest that increases in protein identifications caused by contamination do not necessarily correspond to improved enrichment performance and highlight the utility of the P2 Core Proteome as a complementary quality metric for assessing enrichment quality. Based on the contamination scores and increased protein identifications, we defined three conditions which we used to characterize the effects of contamination.

### Quantitative linearity conserved in P2-based enrichment in presence of platelet contaminated plasma

Using this model, we assessed quantification accuracy by spiking-in 16 heavy isotope labelled recombinant proteins in five two-fold dilution series. Those proteins have all been previously identified in plasma samples and they span a wide range of endogenous expression levels from 8x10^5^ to 1.7x10^10^ pg/L (Table 1). Fourteen of the sixteen proteins were represented by peptides with signals above the calculated LOQ across at least three concentration points. For the majority of proteins, we observed a linear relationship (R^2^ > 0.8) of heavy protein signal and spike-in concentration in at least one contamination condition (Fig. 4A). This result indicates that quantitative information was preserved across a broad dynamic range and that relative abundance changes can be quantified reliably within each of the tested concentration ranges. To assess whether contamination affected quantitative response, we compared concentration-response slopes across contamination backgrounds using a linear mixed-effects model. Most proteins, except ENO2 and GSR exhibited significant Concentration × Contamination interactions (Fig. 4B), indicating that contamination alters the magnitude of the response for specific proteins despite maintained linearity. For CALM1, we could not calculate the calibration curve for the non-contaminated plasma, impeding the proper assessment of the contamination effect. For proteins quantified with more than two peptides above the estimated LOQ, endogenous protein concentrations were estimated using the corresponding light-to-heavy peptide ratios (Fig. 4C). For five out of the 13 quantified proteins, the estimated endogenous levels did not differ between contamination backgrounds, while the remaining eight proteins displayed altered endogenous levels that increased with increase of contamination. Together, these findings indicate that contamination can affect both quantitative protein response and estimated endogenous protein abundance while largely preserving relative quantitative relationships; however, these effects are protein-specific and not necessarily coupled. Additionally, we compared the estimated protein levels to the previously reported protein levels. Apart from the highly abundant APOE, all estimated protein levels either exceeded or fell below the reported concentrations.

**Figure 4.**
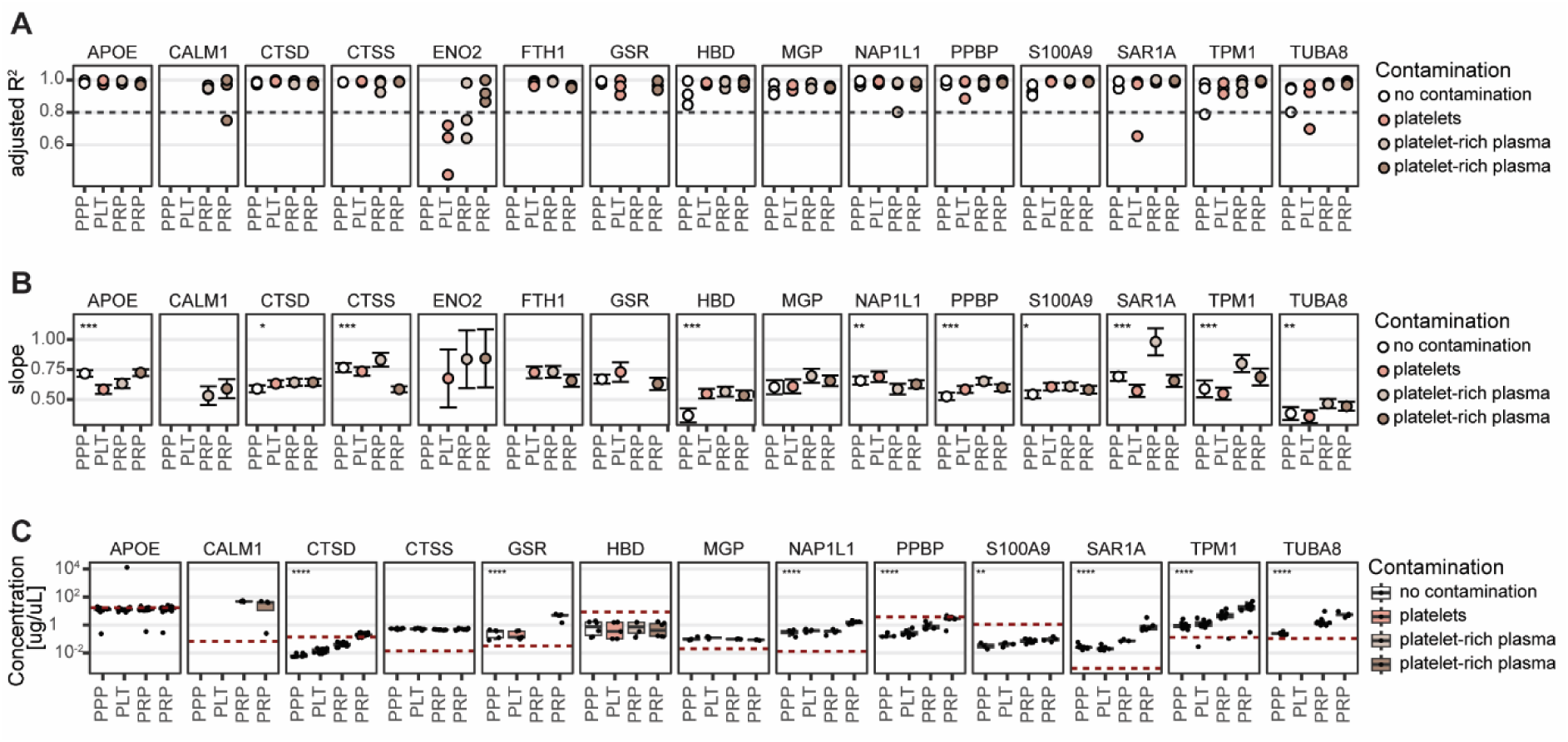
Effects of contamination on quantitative protein response and abundance estimation in P2-enriched plasma. **(A)** Linearity of 15 heavy isotope-labelled recombinant proteins spiked into K_2_EDTA plasma at five two-fold increasing concentration across different contamination backgrounds as defined in Figure S7: uncontaminated platelet-poor plasma (PPP, white), platelet-contaminated plasma (PLT, orange) and plasma supplemented with two levels of platelets-rich plasma (PRP, light and dark brown), followed by P2-based enrichment (n = 3). For each protein, contamination background and technical replicate, linearity between protein quantity and spike-in concentration was assessed by linear regression (see Methods for details) and expressed as the adjusted coefficient of determination (adjusted R^2^). Individual points represent adjusted R^2^ values for each technical replicate. The gray dashed line indicated the predefined threshold for a linear response (adjusted R^2^ = 0.8). **(B)** Slopes of the linear regression models described in (A), representing the change in protein quantity with increasing spike-in concentration across the different contamination backgrounds. Points indicate the estimated regression slopes and error bars indicate 95% confidence intervals. Asterisks indicate significant differences in slope between contamination backgrounds (ANOVA of interaction terms, Benjamini-Hochberg adjusted p-values; *: p < 0.05, **: p < 0.01, ***: p < 0.001, ****: p < 0.0001). **(C)** Estimated endogenous protein concentrations across contamination backgrounds described in (A). Individual points represent endogenous concentration estimates for individual peptides, with their distributions shown as boxplots; mean values represent the estimated endogenous protein concentration. Red dotted lines indicate the corresponding plasma protein concentrations reported in the Human Protein Atlas. Differences in estimated protein concentrations among contamination backgrounds were assessed by ANOVA. Asterisks indicate significant differences (*: p < 0.05, **: p < 0.01, ***: p < 0.001, ****: p < 0.0001).

To assess whether the observed dependence on contamination is intrinsic for the enrichment workflow, we analyzed the same proteins in the same contamination backgrounds using a neat plasma proteomics workflow (Fig. S8). The linear relationship between the heavy protein signal and spike-in concentration was preserved and the data showed low dispersion around the fitted regression line. (Fig. S8B), albeit fewer proteins were overall detected. Consistent with the observations, the contamination affected the quantitative response in neat plasma as well (Fig. S8C), suggesting that the influence of plasma quality on the quantitative response is not an artefact of the enrichment workflow. Interestingly, the estimated endogenous protein levels were mostly stable, only PPBP showed a significant dependence on the contamination background, and the estimated levels increased with increasing contamination. Importantly, for half of the 12 proteins with sufficient peptide-level evidence for protein quantification, abundance estimates could only be obtained in one or two samples, limiting the ability to systematically assess the impact of contamination on endogenous protein levels.

Taken together, these observations demonstrate that contamination preserved the detectability or linear quantification of the spiked proteins but influences the magnitude of the quantitative response and abundance estimates in a protein-specific manner. The comparable effects observed in neat plasma and P2-enriched samples suggest that this dependency is an intrinsic property of plasma quality rather than a consequence of the enrichment workflow itself.

### P2-based enrichment preserves and increases the number of biological signatures while increasing analytical depth in pancreatic cancer pilot study

While the spike-in experiment demonstrated that both neat and P2-enriched plasma workflows preserve linear quantitative relationships across a broad dynamic range, it also revealed that contamination can influence quantitative response and estimated protein abundance in a protein-specific manner. Importantly, comparable contamination-dependent effects were observed in both workflows. Since biomarker discovery relies on the reliable detection of biologically meaningful protein abundance changes, we next investigated two key questions: (i) whether P2-based enrichment preserves disease-associated abundance differences observed in neat plasma and (ii) whether contamination influences the detection of such differences. To address these questions, we compared differential protein abundance patterns between neat and P2-enriched plasma in clinical cohorts and across varying contamination backgrounds.

A cohort comprising K_2_EDTA-plasma specimen from five pancreatic cancer patients and five matched healthy controls was processed both neat and with P2-based enrichment (Fig. 5A). Matching both datasets against the P2 Core Proteome confirmed enrichment quality scores of 93-99.4% (Fig. 3D). P2-based enrichment increased proteome coverage from 958 protein groups identified in neat plasma to 6316 protein groups identified in P2-enriched samples. Moreover, P2 data showed a sharper contrast between pancreatic cancer and healthy control samples in PCA compared to neat data (Fig. 5B). Consistent with the increased analytical depth, P2-based enrichment identified substantially more significantly regulated proteins than neat analysis (797 vs 237 proteins respectively; FDR <5%, |log_2_(fold change)|>0.58) (Fig. 5C). Importantly, 106 significantly regulated proteins were shared between the two workflows of which 97% exhibited the same direction of regulation (Fig. 5D). These proteins included PIGR, a previously reported biomarker for pancreatic cancer^56^. The magnitude of the differential abundance was also highly consistent between workflows, with a strong Pearson correlation (R = 0.71) for the overlapping significantly regulated proteins. Additionally, matching serum samples processed both as neat and P2-enriched samples showed similar results. 92% of 178 overlapping significantly regulated proteins shared directionality with a correlation of fold changes of R = 0.64 (Fig. S9).

**Figure 5:**
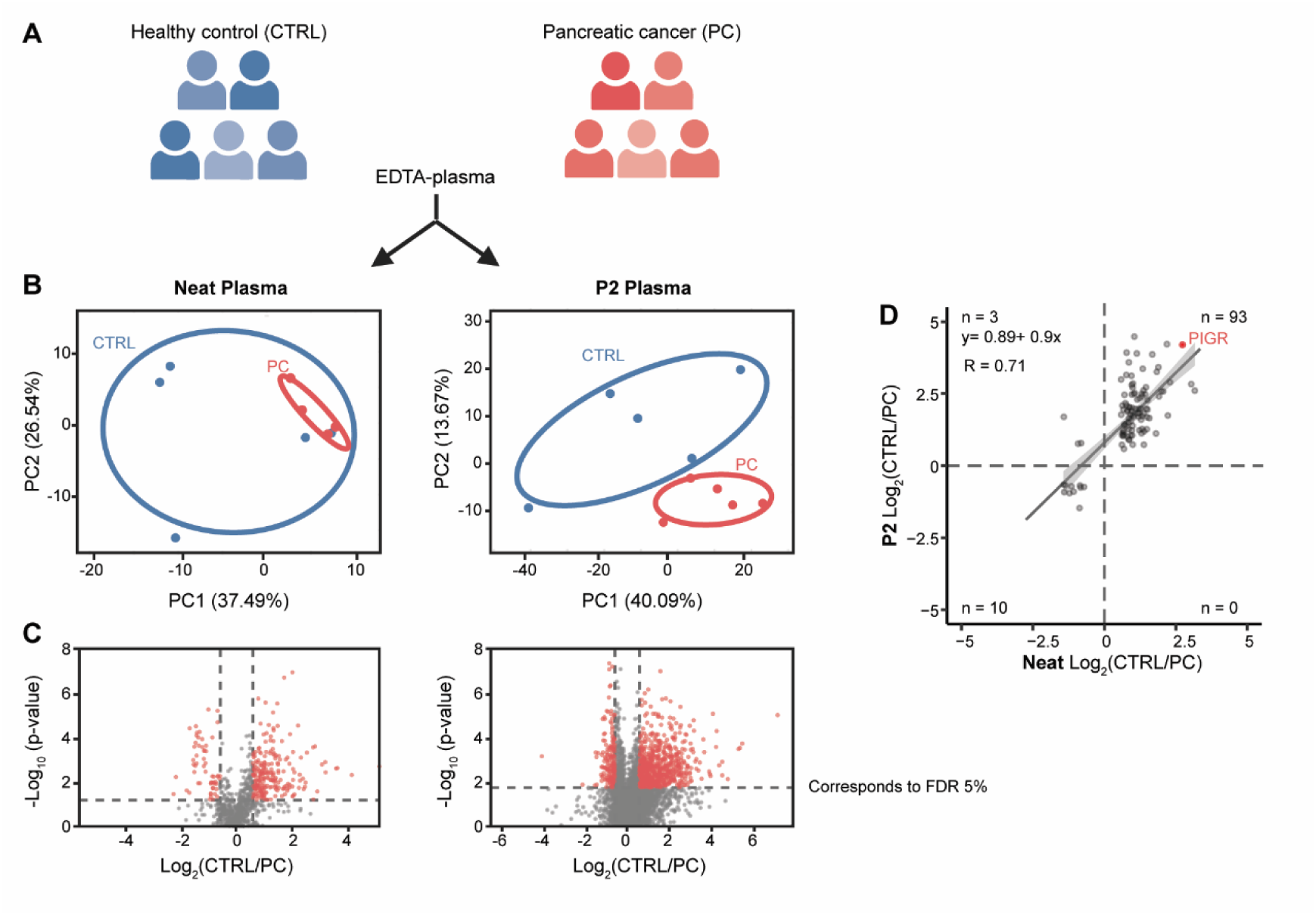
Comparison of disease-associated protein abundance changes in neat and P2-enriched plasma. **(A)** Study design comprising K_2_EDTA-plasma from five individuals with pancreatic cancer (PC) and five healthy controls (CTRL), analyzed as neat plasma and following P2 enrichment. Image generated using BioRender.com (biorender.com). **(B) P**rincipal component analysis (PCA) of protein group quantities in neat plasma (left) and P2-enriched plasma (right). Colors indicate healthy controls (blue) and pancreatic cancer samples (red). **(C)** Differential abundance analysis comparing healthy controls and pancreatic cancer samples in neat plasma (left) and P2-enriched plasma (right, visualized as volcano plots. The log₂ fold change (CTRL/PC) is shown on the x-axis and the −log₁₀(p-value) on the y-axis. Significantly differentially abundant proteins are highlighted in red (FDR < 5% and |log₂ fold change| > 0.58). **(D)** Comparison of log₂ fold changes between healthy controls and pancreatic cancer samples for proteins quantified in neat and P2-enriched plasma. Gray points represent individual proteins, and previously reported biomarkers for pancreatic cancer are highlighted in red. The solid line represents the linear regression between fold changes obtained from the two preparation methods (R = 0.71).

Taken together, these findings demonstrate that P2-based enrichment substantially expands proteome coverage and increases the number of detectable disease-associated proteome changes. P2-based enrichment preserved the biological signatures observed in neat plasma and serum proteomic analyses. The strong consistency of differential abundance patterns between workflows supports the use of P2-based enrichment for biomarker discovery while maintaining compatibility with downstream validation using orthogonal analytical technologies.

### Contamination moderately affects differential abundance testing while preserving disease-associated protein signatures in colorectal cancer study

While our controlled spike-in experiment demonstrated that contamination moderately influences quantitative protein measurements in both neat and P2-based enrichment of plasma (Fig. 4), it remains unclear whether these effects translate into impaired biomarker discovery with biological specimens. Specifically, quantitative changes in individual proteins do not necessarily affect the ability to identify biologically meaningful differences between clinical groups. Hence, we investigated how varying levels of platelet contamination influence statistical power and the detection of differentially abundant proteins in a case-control biomarker discovery setting.

For this, we assembled a second cohort comprising K_2_EDTA-plasma specimens from 12 colorectal cancer patients and 12 matched healthy controls. To establish a controlled contamination model, we first cleared the plasma by centrifugation to remove cellular and debris contamination. The cohort was then processed with P2-based enrichment under two conditions: platelet-poor plasma and plasma supplemented with platelets (Fig 6A). The increase in contamination was confirmed by calculating platelet contamination indices and observed the expected increase for samples with platelet contamination (Fig. S10A and B). Matching the datasets against the P2 Core Proteome confirmed enrichment quality scores of 60.2-93.3% in platelet-poor plasma and 96.6-99% in contaminated plasma (Fig. S10C). To test the influence of contamination, we performed differential analyses across a series of sample combinations containing increasing proportions of contaminated samples (Fig. 6B). We ensured that the contaminated samples were distributed equally between the healthy and colorectal cancer groups in combinations a to f, thereby modelling increasing plasma variability without confounding disease status. In contrast, combination g intentionally confounded contamination within disease status by assigning cleared samples to the healthy group and contaminated samples to the colorectal cancer group.

**Figure 6:**
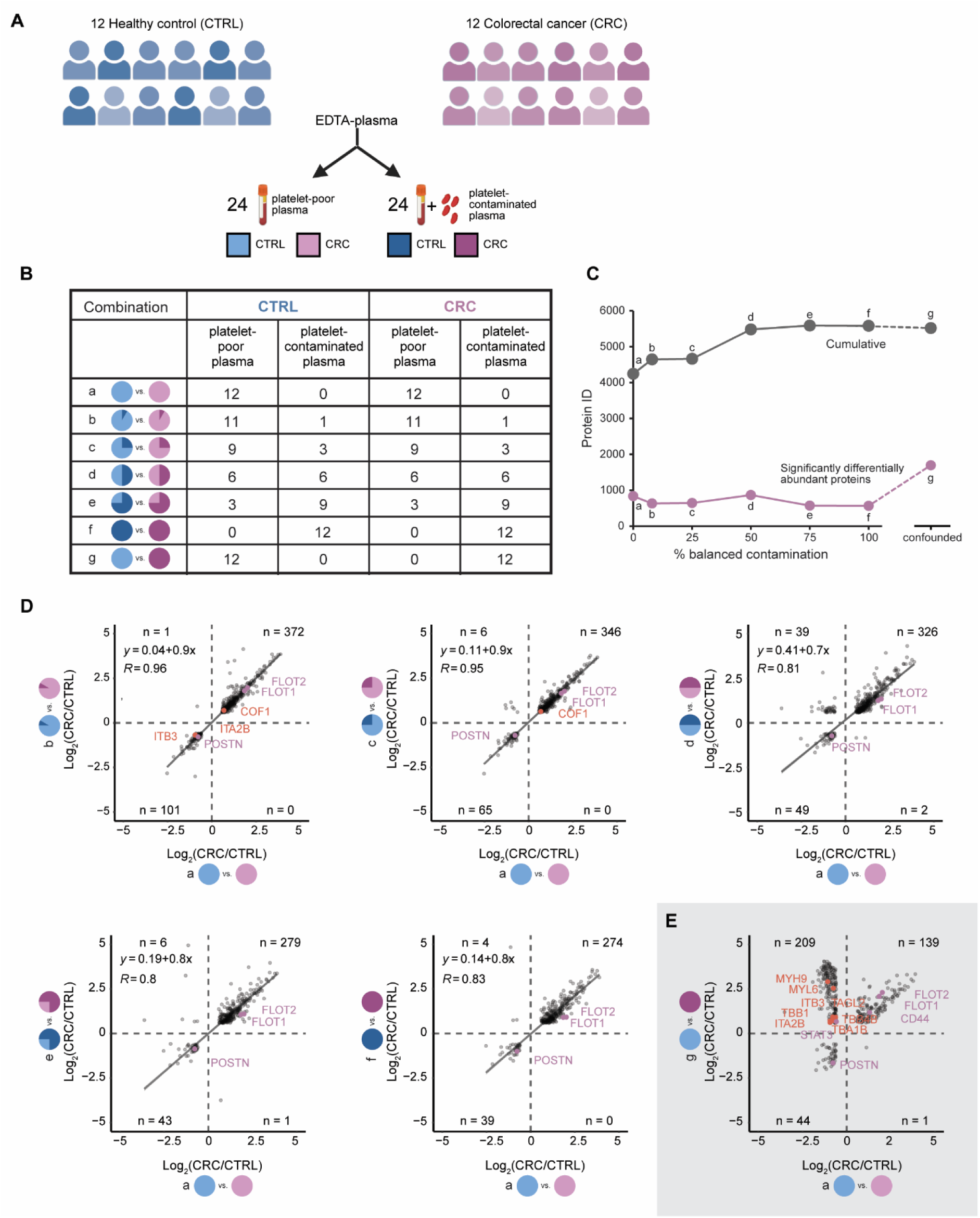
Effect of platelet contamination on disease-associated protein abundance changes in P2-enriched plasma. (**A**) Study design comprising platelet-poor K₂EDTA plasma from 12 individuals with colorectal cancer (CRC; purple) and 12 healthy controls (CTRL; blue). Each sample was processed either without contamination (lighter shades) or with controlled platelet contamination (darker shades) and subsequent P2 enrichment. **(B)** Schematic representation of analysis combinations a-g used for differential abundance analysis, comprising different proportions of platelet-poor and platelet-contaminated samples. In combinations b-f, platelet contamination is introduced symmetrically between CTRL and CRC groups, the combination g represents asymmetric contamination, with platelet-poor CTRL samples compared with platelet-contaminated CRC samples. **(C)** Cumulative number of protein group identifications (gray) and significantly differentially abundant proteins between CTRL and CRC (purple) for combinations a–f as a function of the proportion of platelet-contaminated samples and the confounded sample analysis with combination g. **(D, E)** Comparison of log_2_ fold changes (CRC/CTRL) obtained from uncontaminated samples (combination a; a-axis) with those obtained from (D) combinations b-f with increasing symmetric platelet contamination or (E) combination g with asymmetric platelet contamination. Gray points represent individual proteins, selected colorectal cancer-associated proteins are highlighted in pink, platelet-associated proteins in orange. Solid lines represent linear regression fits between log_2_ fold changes, with the corresponding regression equation and Pearson correlation coefficient (R), indicated in each panel.

Despite the increase in analytical depth introduced by platelet contamination (from 4244 to 5580 protein groups), the number of significantly differentially abundant proteins remained largely stable across the different contamination scenarios, ranging from 564 to 834 proteins (Fig. 6C). Importantly, this set included established colorectal cancer biomarker candidates, such as CD44, FLOT1, FLOT2, POSTN and STAT3^57–61^. PCA showed similar separation of health and disease in combinations a to f (Fig. S11).

Next, we compared the differential abundance results obtained from the contamination scenarios (combination b to f) with those from the clean spun plasma reference (combination a). The majority of shared significantly regulated proteins showed highly consistent effect sizes, with Pearson correlation coefficients decreasing only moderately from R=0.9 to R=0.8 as the proportion of contaminated samples increased (Fig. 6D). Increasing contamination introduced a systematic attenuation of fold changes, reflected by a decrease in regression slope from 0.96 to 0.80. The direction of differential abundance remained preserved for the majority of the proteins, including the established colorectal cancer biomarkers. These findings indicate that platelet contamination modestly reduces statistical contrast rather than fundamentally altering the biological conclusions.

To model a worst-case scenario that can be encountered in clinical studies, we generated the confounded dataset (combination g), in which all healthy control samples consisted of platelet-poor plasma, while all colorectal cancer samples contained platelet contamination. Under these conditions, the number of significantly differentially abundant proteins in the confounded dataset increased substantially to 1702, nearly twice as many as in the balanced platelet-poor comparison. Many of these proteins corresponded to known platelet-associated markers, including MYH9, MYL6, ITB3, ITA2B, TAGL2, TBR4B, TBA1B, TBB1^34^ (Fig. 6E). Comparison of differential abundance results between the balanced and confounded dataset revealed systematically larger fold changes in the confounded condition. Importantly, despite this inflation of effect sizes, the direction of differential abundance was preserved for all except one of the detected CRC biomarkers.

## Discussion

Particle-based enrichment workflows have enabled LC-MS-based plasma proteomics to move beyond the several hundred proteins typically detected in neat plasma or serum. Recent studies have highlighted their sensitivity to pre-analytical variation^23,33,34^. To advance the understanding of particle-based enrichment workflows, we systematically characterized the P2-based enrichment workflow under real-world conditions and across a wide range of commonly encountered pre-analytical variables. To our knowledge, this study represents the most comprehensive characterization of enrichment workflows across preanalytical variables. We showed that proteome composition and depth were affected by collection matrix, sample handling, pathogen inactivation procedures, and cellular contamination, although the magnitude of these effects varied substantially between variables. These insights enable selection of the most appropriate study design and allow estimation of the impact of specific preanalytical factors.

A recurring question surrounding enrichment technologies is whether increased proteome depth reflects meaningful biology or mostly increased detection of preanalytical artefacts. Our matched analysis of neat and P2-enriched plasma directly addresses this concern. Despite a substantial increase in proteome coverage, 97% of overlapping significantly regulated proteins showed the same direction of regulation between the two workflows, with similar findings observed in serum. These results indicate that P2 enrichment largely preserves disease-associated biological information while expanding access to additional proteins. Our findings are consistent with recent studies showing that platelet, erythrocyte, and coagulation-related contamination can substantially affect nanoparticle-based plasma proteomics workflows.^23,33,34^ These studies demonstrated that cellular contamination increases proteome depth by introducing blood cell-derived proteins and can lead to preanalytical bias in biomarker discovery. In agreement with these reports, platelet contamination increased protein identifications following P2 enrichment, whereas erythrocyte contamination produced more moderate effects. Importantly, our study extends previous work by showing that cellular contamination does not necessarily compromise biological interpretation. When the cellular contamination was distributed evenly between study groups, disease-associated signatures and fold-change directionality were largely preserved. In contrast, when the cellular contamination was confounded with disease status, effect sizes were inflated and numerous additional significant proteins appeared, many of which represented known platelet-associated markers. It is worth noting that these findings were derived from pilot-scale studies in which cellular contamination was introduced after blood collection. Further studies using prospectively collected clinical samples will be valuable to confirm the observed effects. These findings provide experimental evidence that preanalytical variables can generate apparent disease signatures when they are confounded with biology. Notably, the colorectal cancer contamination experiment shows that greater analytical depth alone does not guarantee improved biomarker discovery. Still, the quality of biological conclusions remains dependent on controlling preanalytical variation and avoiding confounding between sample quality and disease status. Beyond the biological implications of cellular contamination, an additional question is whether P2-based enrichment preserves quantitative accuracy for protein measurements. Building on previous work demonstrating absolute quantification using heavy-labeled proteins spiked into neat plasma^50^, we evaluated the quantification of recombinant heavy-labeled proteins in platelet-poor plasma and plasma samples with cellular contamination. Relative quantification remained highly reproducible and comparable to that observed in neat plasma. Moreover, quantitative measurements showed high precision across workflow replicates. In contrast, absolute protein concentrations estimated after P2-based enrichment differed from those obtained in neat plasma and from concentrations reported in the Human Plasma Protein Atlas^62^. This discrepancy is likely explained by the recombinant nature of the spiked proteins, which lack endogenous post-translational modifications, physiological interaction partners, and other native molecular features that may affect enrichment efficiency. These observations warrant further systematic investigation.

The observation of a highly reproducible protein set motivated us to establish the P2 Core Proteome. It can be viewed as a mass spectrometry-based analogue of targeted affinity-proteomics panels such as those employed by Olink, Alamar Biosciences or SomaLogic. The large overlap with the Human Plasma Protein Atlas and extracellular vesicle-associated proteins suggests that this panel reflects proteins consistently accessible by P2-based enrichment. Still, it should not be interpreted as a comprehensive representation of the P2 plasma proteome, as it is likely lacking disease-specific proteins. Instead, its main value lies in serving as a robust quality-control and benchmarking metric for evaluating enrichment performance across studies and laboratories. Further, we plan to explore its use for normalization and batch correction of large studies.

Preanalytical variables are an inherent component of plasma biomarker discovery studies. Building on previous largely qualitative reports^23,33,34^ and the quantitative analyses presented here, we conclude that P2 and related particle-based enrichment technologies provide unique access to deep plasma proteomes. A key finding of this study is that biological signals identified through P2-based discovery proteomics appear to be highly reproducible in downstream targeted assays, including SISCAPA- and affinity-based approaches. These findings support the use of P2-derived biomarker candidates in bioanalytical validation workflows aligned with ICH M10 and FDA guidelines. This may facilitate the translation of deep-discovery proteomics findings into regulated biomarker development workflows.

## Author Contributions

A.V., L.M., Y.F., A.L., N.K., O.R and R.B. designed the project. A.V., L.M., P.S., S.S., Y.F., L.R., I.A. and R.B. supported the experimental design of the research. A.V., L.M., S.S., V.B., K.K., N.B., C.R.B., W.V.B. prepared samples and carried out the measurements. A.V., P.S., S.S., L.M. and R.B. performed the data analysis. A.V., L.M., P.S. and R.B. wrote the paper. R.B. supervised the project. All authors critically revised the manuscript and approved its content.

## Data availability

The raw MS data, the spectral libraries and the quantitative data tables have been deposited to the ProteomeXchange Consortium via the PRIDE partner repository with the dataset identifier XXXX. The Saved projects from Spectronaut can be viewed with the Spectronaut Viewer (www.biognosys.com/spectronaut-viewer).

## Acknowledgements

We would like to thank Martin Menert, Katharina Limm and P.Ittu for input to the manuscript. We thank Sebastian Müller and Kevin Pacios for support during sample acquisition and Philip Lössl and Phoebe Wei for guidance and support with the recombinant, heavy labelled proteins. We would like to dedicate this work to Lukas Reiter for his contribution that helped shaping this work.

## Abbreviations

ACD: Acid Citrate Dextrose
ANOVA: Analysis of Variance
CPD: Citrate Phosphate Dextrose
CRC: Colorectal Cancer
CV: Coefficient of Variation
DIA: Data-Independent Acquisition
EDTA: Ethylenediaminetetraacetic Acid
EV: Extracellular Vesicle
FDR: False Discovery Rate
GO: Gene Ontology
HLB: Hydrophilic-Lipophilic Balanced
LC-MS: Liquid Chromatography-Mass Spectrometry
LOQ: Limit of Quantification
PAC: Protein Aggregation Capture
PCA: Principal Component Analysis
PC: Pancreatic Cancer
PBS: Phosphate-Buffered Saline
PBMC: Peripheral Blood Mononuclear Cell
P2: Plasma Enrichment Technology P2
PPP: Platelet-Poor Plasma
PRP: Platelet-Rich Plasma
QC: Quality Control
R: Coefficient of Determination
RBC: Red Blood Cell

---

**Figure S1:**
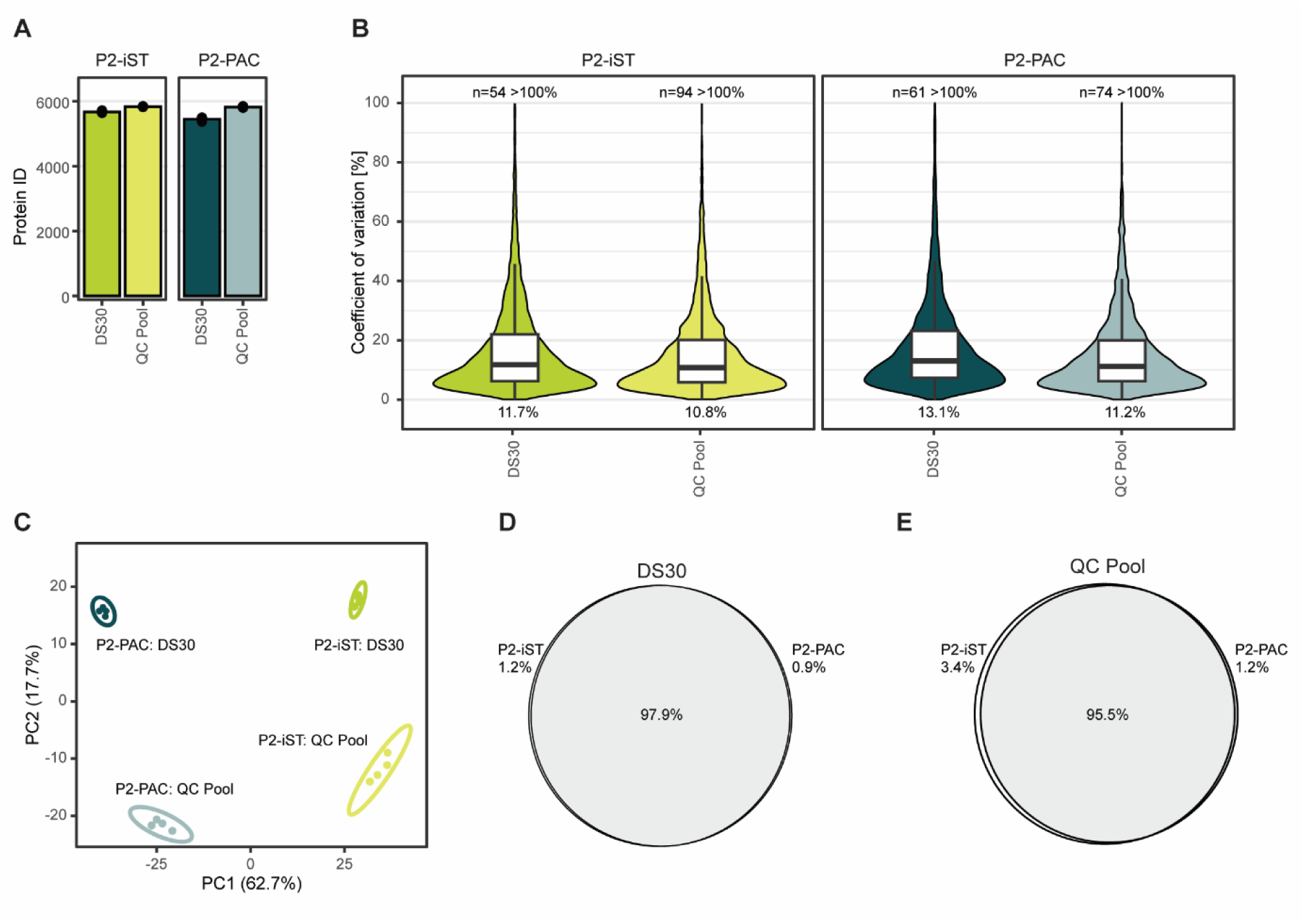
P2-PAC and P2-iST Plasma kit comparison. K₂EDTA-plasma pools (named DS30 and QC) were prepared in quadruplicates with P2-PAC and P2-iST Plasma and acquires shuffle to compare the performance. (**A**) Protein group identifications were counted (**B**), and coefficients of variation were calculated for the quadruplicates. (**C**) Principal component analysis was performed for the two P2-based enrichment workflows. (**D**) VENN diagram of the protein group analyses of the K_2_EDTA-plasma pool DS30 (**E**) VENN diagram of the protein group analyses of the K_2_EDTA-plasma QC pool.

**Figure S2:**
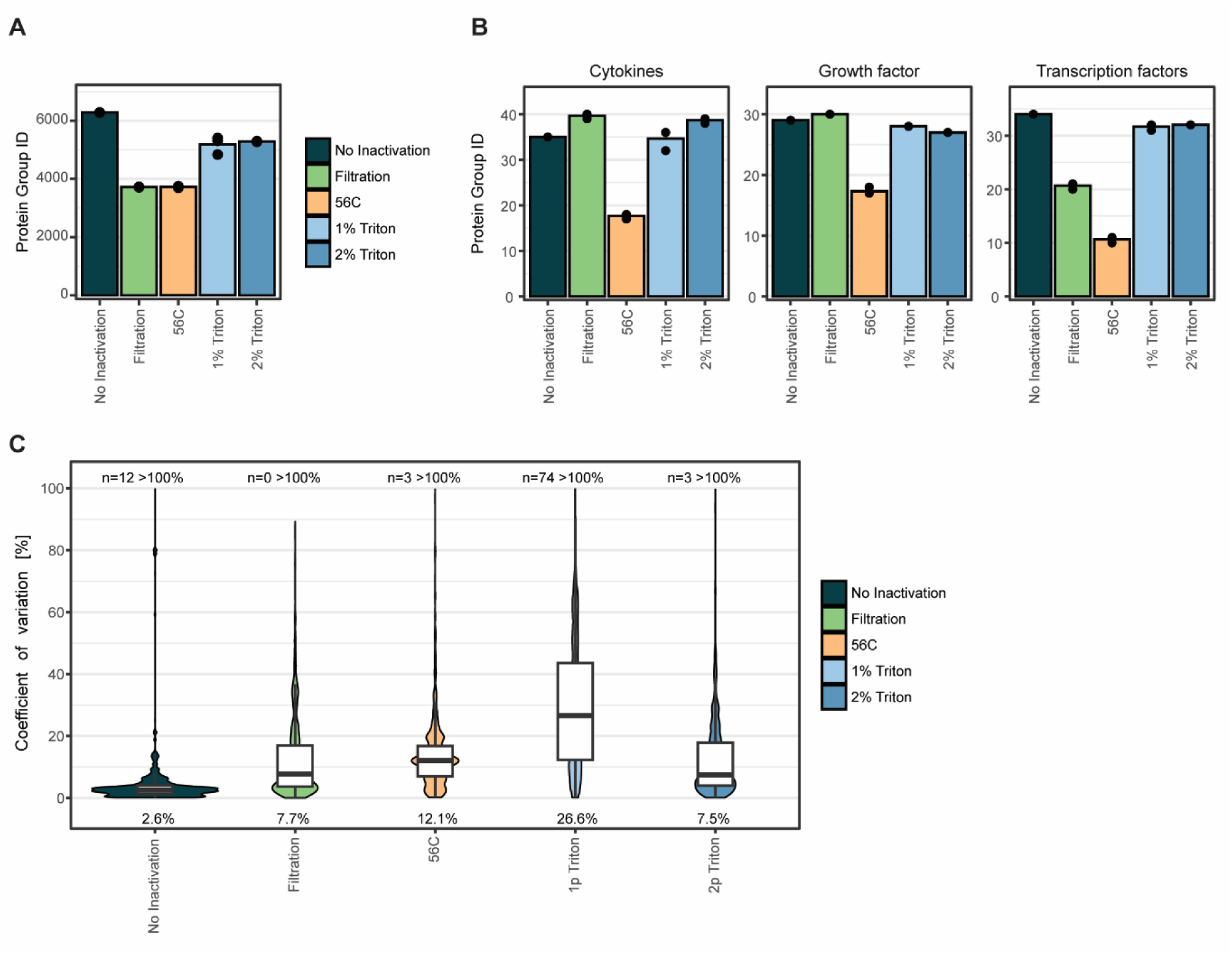
Effect of plasma inactivation methods on P2-based enrichment performance. **(A)** Comparison of protein group identifications following P2-based enrichment of K_2_EDTA -plasma QC pool without inactivation pre-treatment (dark teal) or following inactivation by filtration through 0.22 µ m filters (green), heat treatment at 56 °C for 1 h (orange) or treatment with 1% or 2% Triton-X100 (light blue and dark blue, respectively) (n=3). Mean protein group identifications are displayed as bars, with individual measurements shown as dots. **(B)** Protein identification for selected protein classes (Cytokines, Growth factors, Transcription factors) in the samples described in (A). **(C)** Coefficients of variation (CVs) calculated at the protein group level for the samples described in (A) and displayed as violin plots with overlaid boxplot. Mean CV values are indicated at the bottom. Outliers (>1.5 x interquartile range) are not displayed. CVs >100% are excluded; the number of excluded observations is indicated at the top.

**Figure S3:**
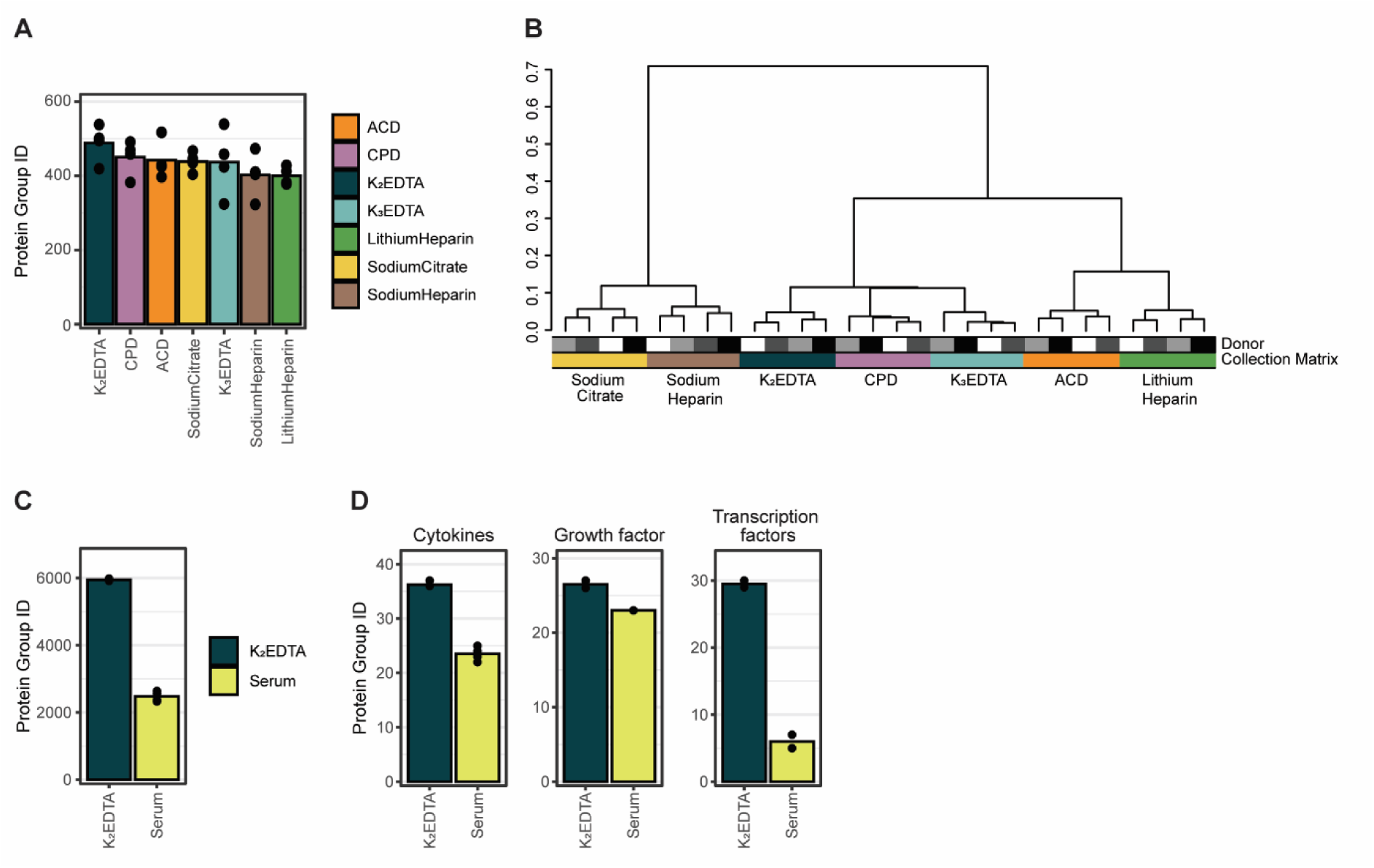
Effect of blood collection matrices and blood components on P2-based enrichment performance. (**A**) Comparison of protein group identifications in neat plasma collected from four donors using different blood collection matrices (dark teal: K_2_EDTA; light teal: K_3_EDTA; purple, CPD; yellow, sodium-citrate; orange, ACD; green, lithium-heparin; brown, sodium-heparin). Mean protein group identifications are displayed as bars, with individual measurements shown as dots. **(B)** Hierarchical clustering analysis based on protein group quantities of the samples described in Figure 2 using Pearson correlation coefficient as distance metric. (**C** and **D**) Comparison of total protein group identifications (C) and protein group identifications within selected protein classes (D) in K_2_EDTA plasma (dark teal) or serum (lime yellow) following P2 enrichment. Mean protein group identifications are displayed as bars, with individual measurements shown as dots.

**Figure S4:**
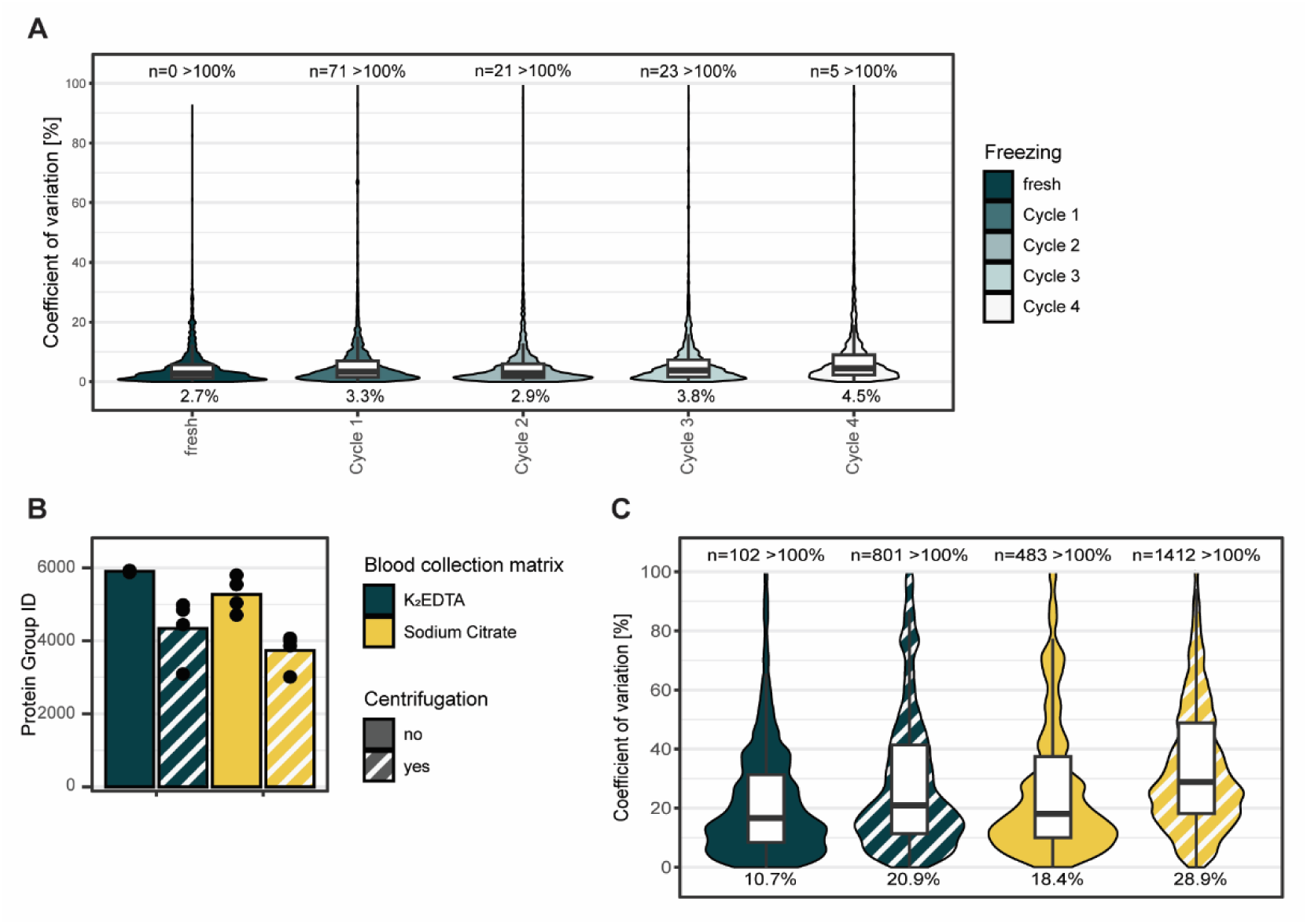
Impact of freeze thaw cycles and centrifugation on the P2-based enrichment workflow. **(A)** Coefficients of variation (CVs) calculated at the protein group level for K_2_EDTA plasma subjected to zero to four consecutive freeze-thaw cycles (dark teal to white). **(B)** Comparison of protein group identifications in K_2_EDTA (dark teal) and sodium-citrate (yellow) plasma from four donors before and after an additional centrifugation step at 2,000 x g (striped bar). Mean protein group identifications are displayed as bars, with individual measurements shown as dots. **(C)** Coefficients of variation (CVs) of the four donors were calculated at the protein group level for the samples described in (B). In (A) and (C), CVs are displayed as violin plots with overlaid boxplots, with mean CV values indicated at the bottom. Outliers (>1.5 x interquartile range) are not displayed. CVs >100% are excluded; the number of excluded observations is indicated at the top

**Figure S5:**
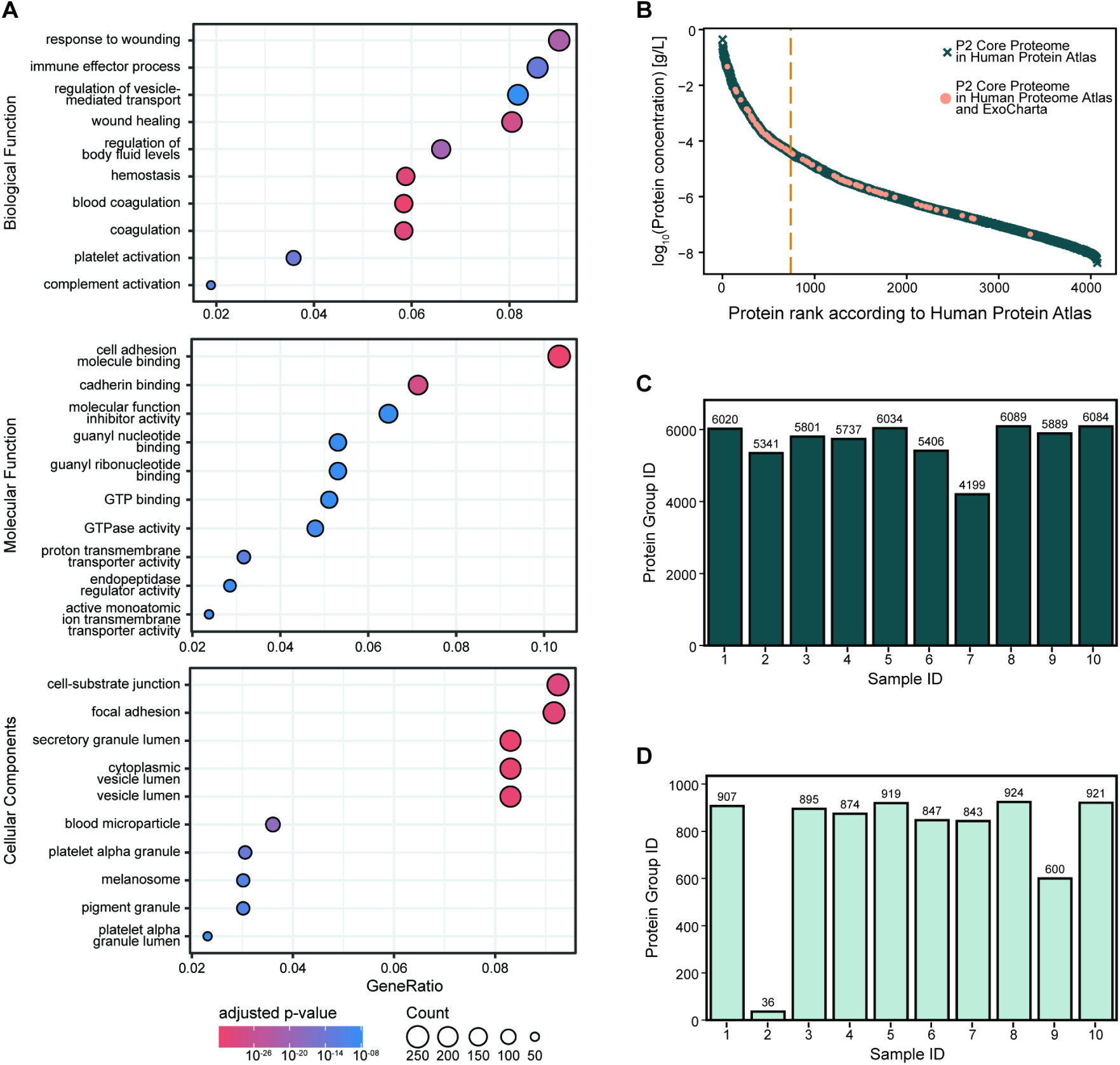
Characterization of the P2 Core Proteome. **(A)** Gene Ontology (GO) overrepresentation analysis of P2 Core Proteome proteins for biological processes (BP, top), molecular functions (MF, middle), and cellular components (CC, bottom). GeneRatio is shown on the x-axis, circle size represents the number of proteins associated with each GO term, and color indicates the adjusted p-value. **(B)** P2 Core Proteome proteins ranked according to protein concentration reported in the Human Protein Atlas (HPA). Proteins also reported in ExoCarta^54^ are highlighted in orange. Dashed orange line represents the median rank of P2 Core Proteins in the Human Protein and ExoCarta (rank = 741.0) **(C, D)** Comparison of protein group identifications following P2 enrichment (C) and in neat plasma (D) across 10 independent samples from the pancreatic cancer cohort.

**Figure S6:**
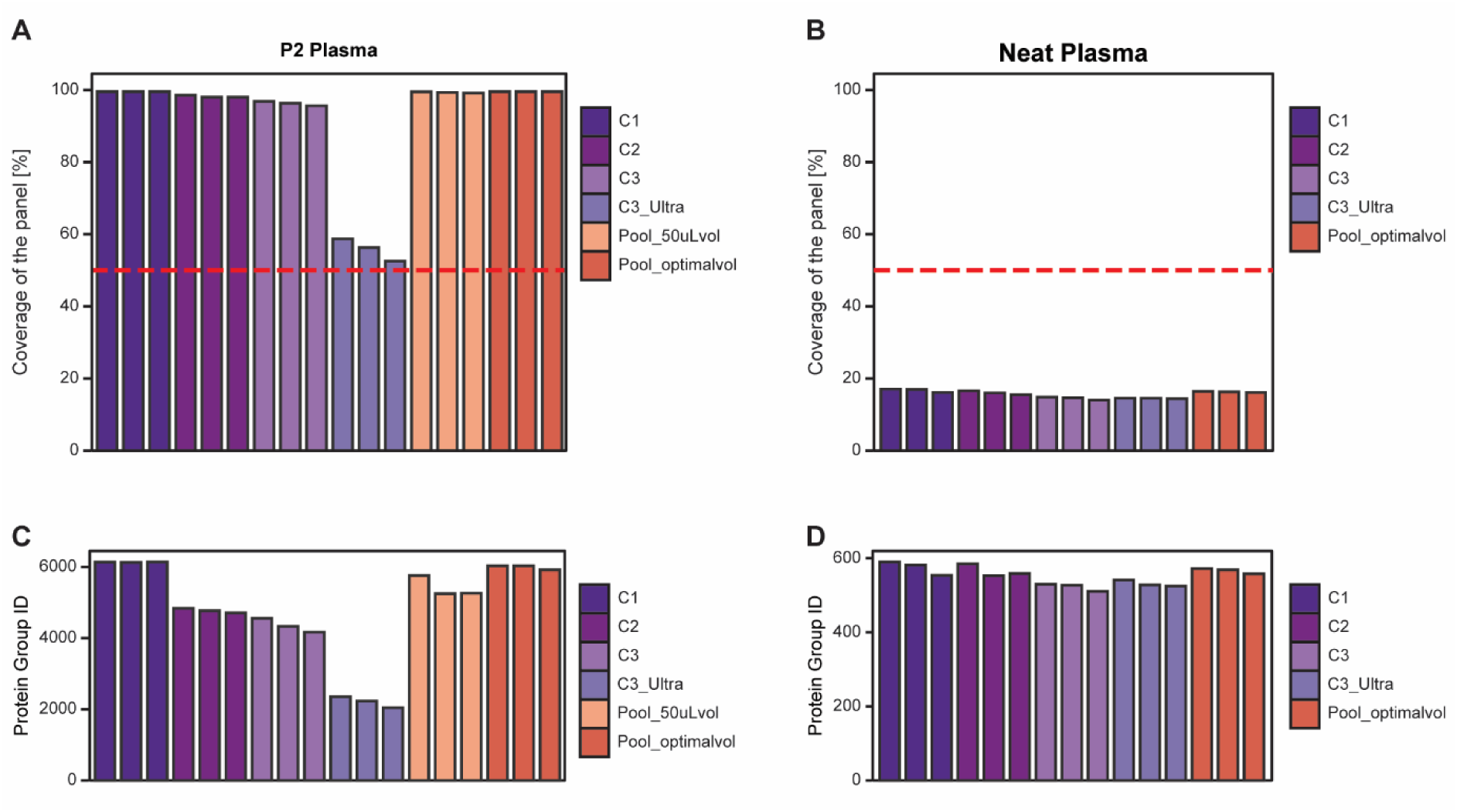
Assessment of generalizability of the enrichment performance score. (**A, B**) P2 Core Proteome coverage in (A) P2-enriched EDTA plasma and (B) neat plasma in the study published by Völlmy et al.^23^, encompassing different pre-analytical conditions, including ultracentrifugation. Different experimental conditions, as described in the study, are indicated by color. Red dashed line indicates the predefined quality threshold of 50% P2 Core Proteome coverage. (**C, D**) Corresponding protein group identifications in (C) P2-enriched plasma and (D) neat plasma for the samples shown in (A) and (B), respectively.

**Figure S7:**
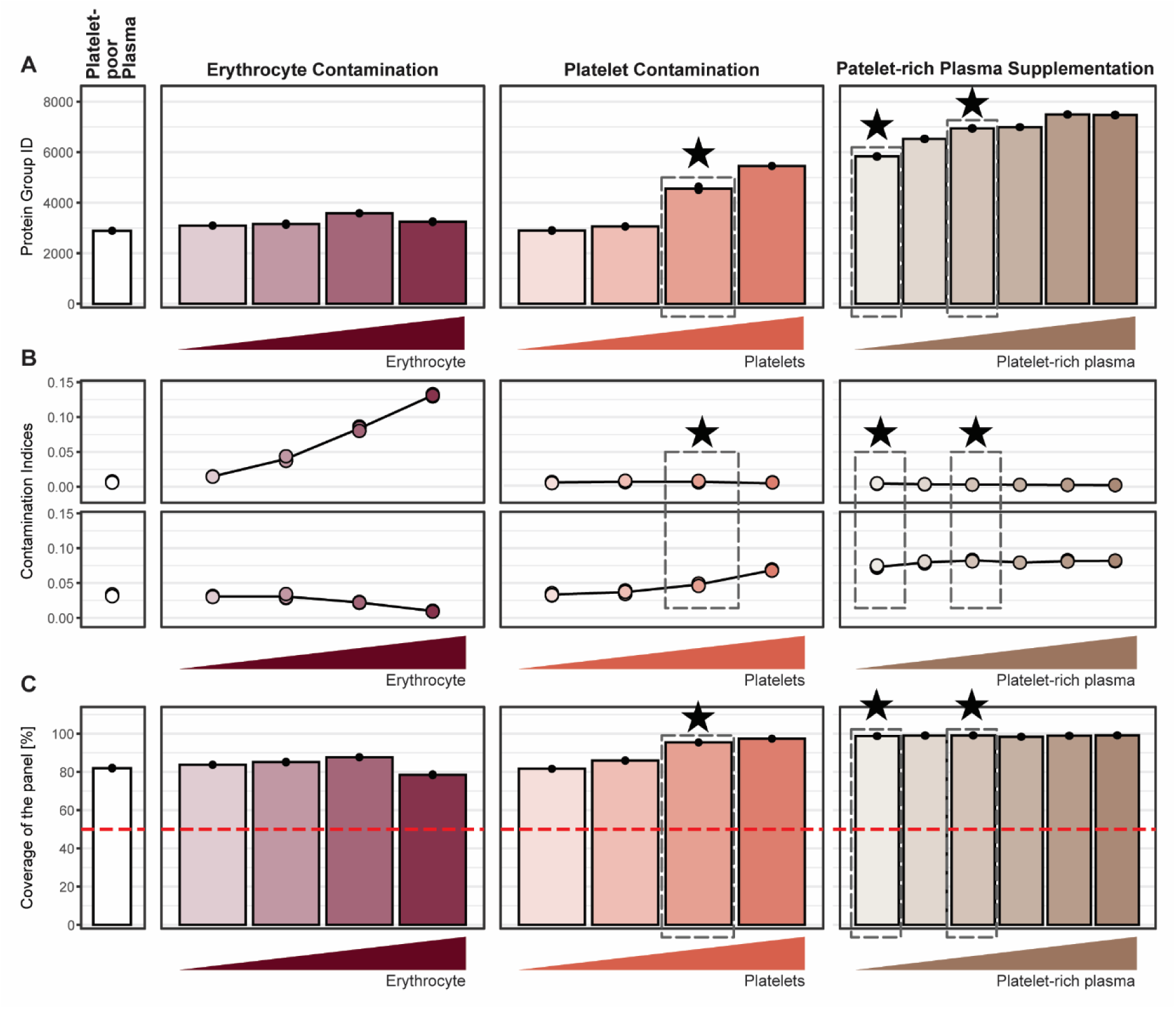
Characterization of the controlled spike-in contamination model. **(A)** Comparison of protein group identifications in platelet-poor plasma (white) and plasma with increasing levels of erythrocyte contamination (red), platelet contamination (orange), or platelet-rich plasma supplementation (brown) (see Method for details). **(B)** erythrocyte (upper row) and platelet (lower row) contamination indices, as defined by Korff et al.^34^ for the samples described in (A).**(C)** P2 Core Proteome coverage in the samples described in (A). Red dashed line indicates the predefined quality threshold of 50% P2 Core Proteome coverage. In (A-C), dashed boxes and stars indicate the contamination conditions selected for further analysis.

**Figure S8:**
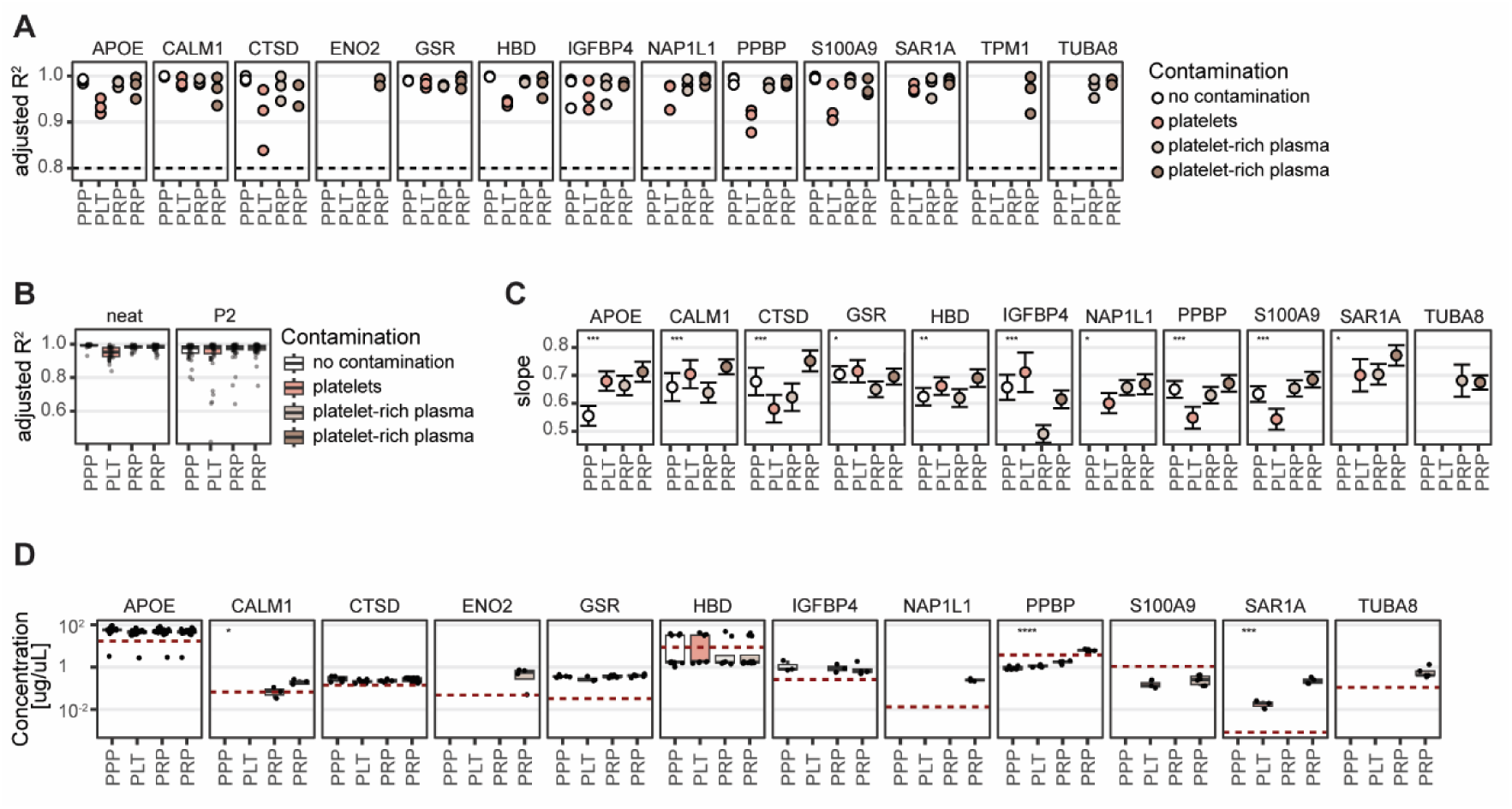
Effects of contamination on quantitative protein response and abundance estimation in plasma. **(A)** Linearity of 15 heavy isotope-labeled recombinant proteins spiked into K₂EDTA plasma at five two-fold increasing concentrations across different contamination backgrounds as defined in Figure S7: uncontaminated platelet-poor plasma (PPP; white), platelet-contaminated plasma (PLT; orange), and plasma supplemented with two levels of platelet-rich plasma (PRP; light and dark brown) (n = 3). For each protein, contamination background, and technical replicate, linearity between protein quantity and spike-in concentration was assessed by linear regression (see Methods) and expressed as the adjusted coefficient of determination (adjusted R²). Individual points represent adjusted R² values for each technical replicate. The gray dashed line indicates the predefined threshold for a linear response (adjusted R² = 0.8). **(B)** Distribution of adjusted R² values across all proteins and contamination backgrounds for neat and P2-enriched plasma. Individual points represent adjusted R² values obtained for each protein and technical replicate under the respective contamination background, with distributions summarized as boxplots. The gray dashed line indicates the predefined threshold for a linear response (adjusted R² = 0.8). **(C)** Slopes of the linear regression models described in (A) representing the change in protein quantity with increasing spike-in concentration across the different contamination backgrounds. Points indicate the estimated regression slopes and error bars indicate 95% confidence intervals. Asterisks indicate significant differences in slope between contamination backgrounds (ANOVA of interaction terms, Benjamini-Hochberg adjusted p-values; *: p < 0.05, **: p < 0.01, ***: p < 0.001, ****: p < 0.0001). **(D)** Estimated endogenous protein concentrations across contamination backgrounds described in (A). Individual points represent endogenous concentration estimates for individual peptides, with their distributions shown as boxplots; mean values represent the estimated endogenous protein concentration. Red dotted lines indicate the corresponding plasma protein concentrations reported in the Human Protein Atlas. Differences in estimated protein concentrations among contamination backgrounds were assessed by ANOVA. Asterisks indicate significant differences (*: p < 0.05, **: p < 0.01, ***: p < 0.001, ****: p < 0.0001).

**Figure S9:**
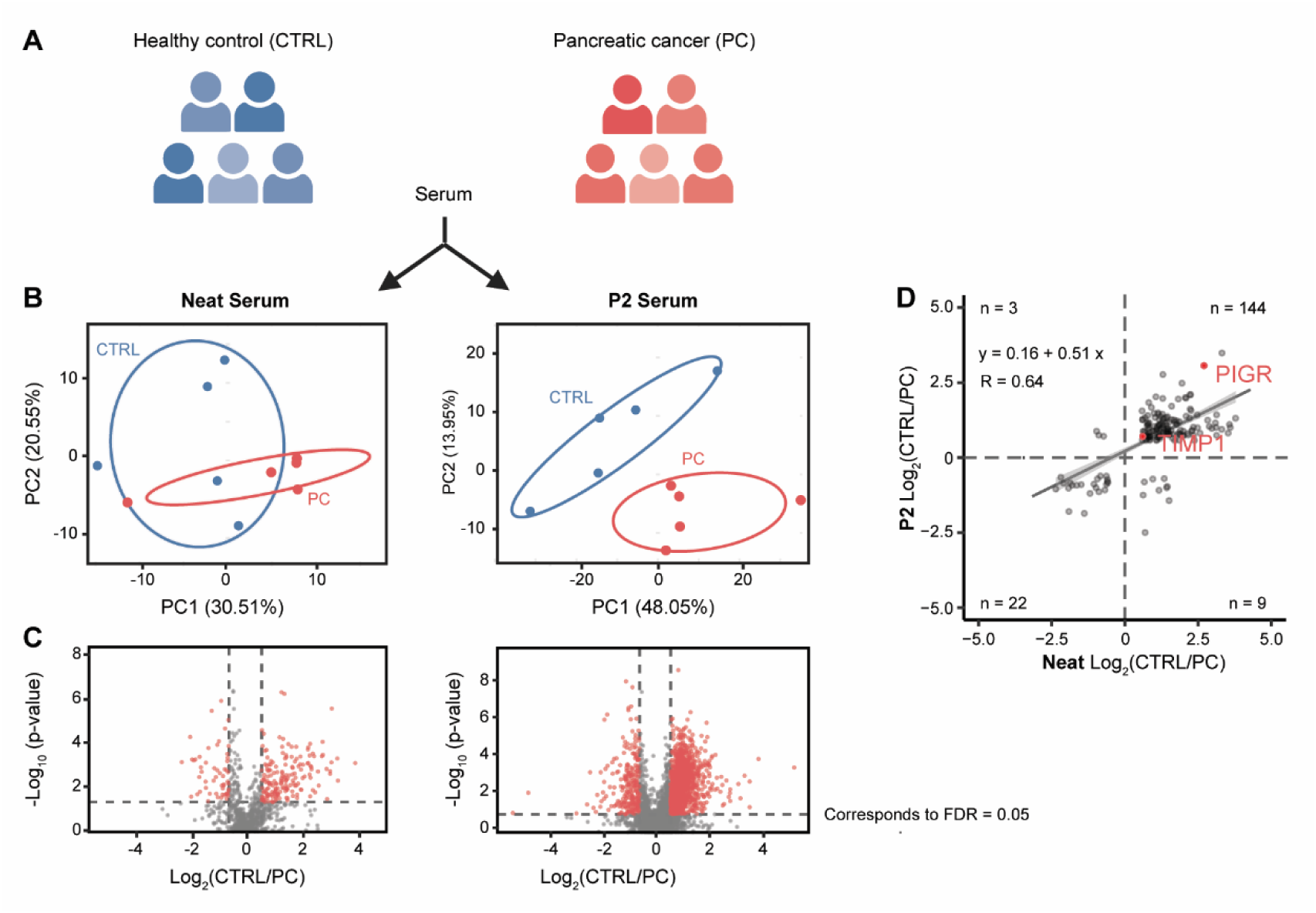
Comparison of disease-associated protein abundance changes in neat and P2-enriched serum. **(A)** Study design comprising serum from five individuals with pancreatic cancer (PC) and five healthy controls (CTRL) as described in Figure 5, analyzed as neat plasma and following P2 enrichment. Image generated using BioRender.com (biorender.com). **(B)** Principal component analysis (PCA) of protein group quantities in neat plasma (left) and P2-enriched plasma (right). Colors indicate healthy controls (blue) and pancreatic cancer samples (red). **(C)** Differential abundance analysis comparing healthy controls and pancreatic cancer samples in neat plasma (left) and P2-enriched plasma (right, visualized as volcano plots. The log₂ fold change (CTRL/PC) is shown on the x-axis and the −log₁₀(p-value) on the y-axis. Significantly differentially abundant proteins are highlighted in red (FDR < 5% and |log₂ fold change| > 0.58). **(D)** Comparison of log₂ fold changes between healthy controls and pancreatic cancer samples for proteins quantified in neat and P2-enriched plasma. Gray points represent individual proteins, and previously reported biomarkers for pancreatic cancer are highlighted in red. The solid line represents the linear regression between fold changes obtained from the two preparation methods (R = 0.64).

**Figure S10:**
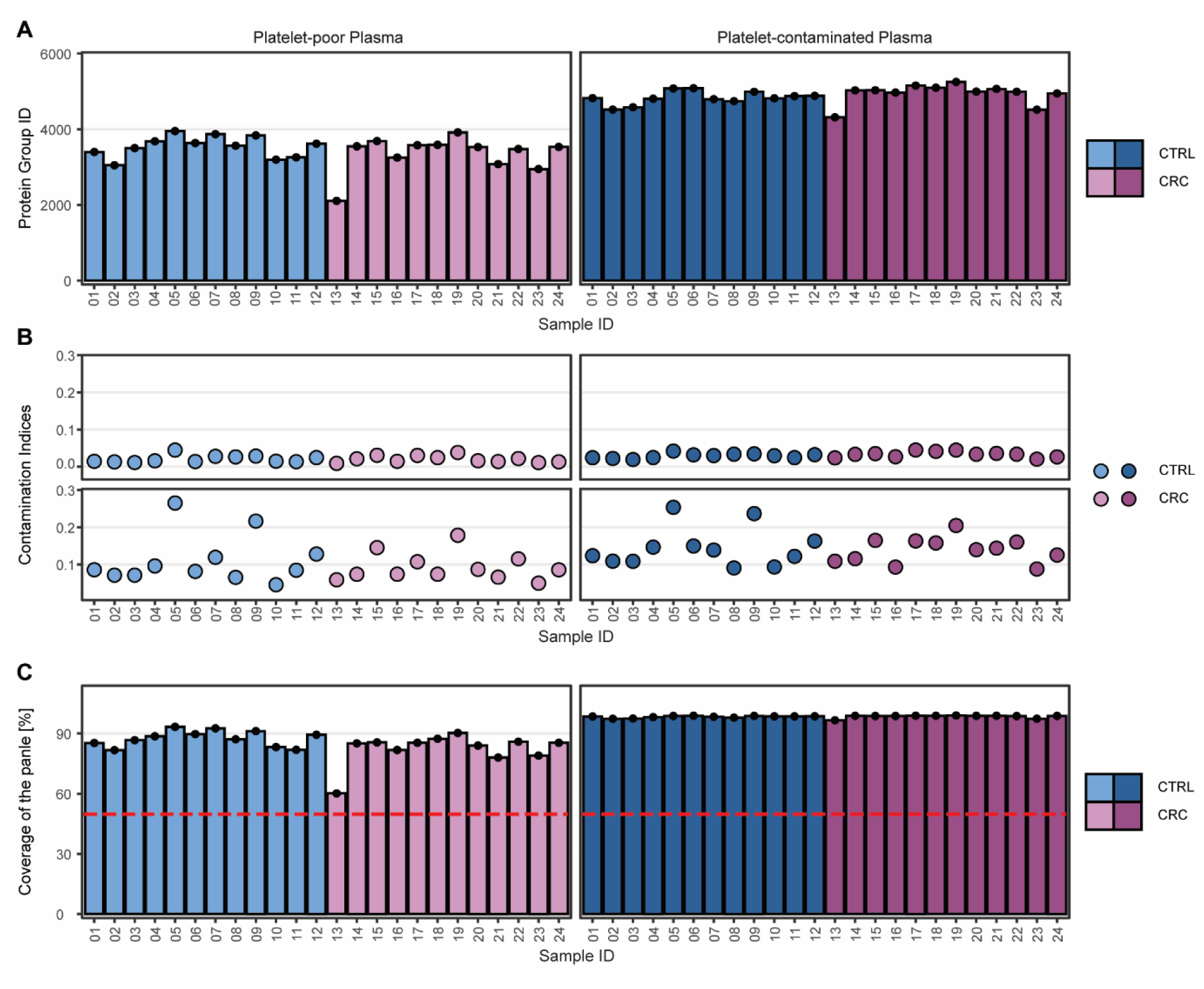
Characterization of the controlled spike-in colorectal cancer cohort. **(A)** Comparison of protein group identifications in 12 colorectal cancer (CRC; purple) and 12 healthy controls (CTRL; blue) without contamination (lighter shades) or with controlled platelet contamination (darker shades). **(B)** Erythrocyte (upper row) and platelet (lower row) contamination indices, as defined by Korff et al.^34^ for the samples described in (A).**(C)** P2 Core Proteome coverage in the samples described in (A). Red dashed line indicates the predefined quality threshold of 50% P2 Core Proteome coverage.

**Figure S11:**
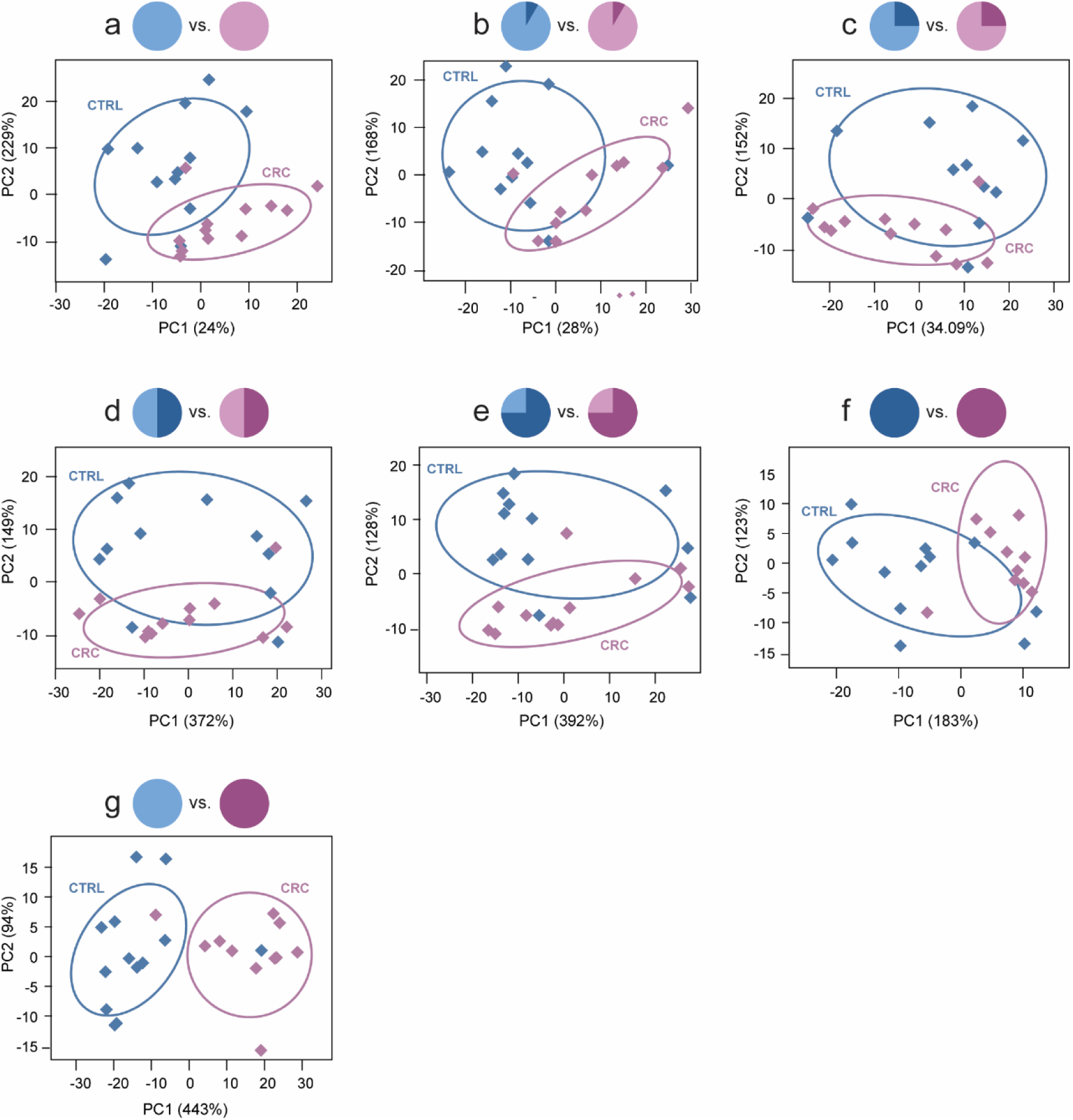
Characterization analysis combinations (a-g) Principal component analyses of the combinations a-g, comprising different proportions of platelet-poor and platelet-contaminated samples. Platelet-poor K_2_EDTA-plasma from 12 individuals with colorectal cancer (CRC; purple) and 12 healthy controls (CTRL; blue).

## Bibliography

1. Cooper, G. M. The Cell. Sinauer Associates https://www.ncbi.nlm.nih.gov/books/NBK9839/ (2000).

2. Alberts, B. et al. Protein Function. in Molecular Biology of the Cell. 4th edition (Garland Science, 2002).

3. Labbadia, J. & Morimoto, R. I. The biology of proteostasis in aging and disease. Annu. Rev. Biochem. 84, 435–464 (2015).

4. Anderson, N. L. & Anderson, N. G. The human plasma proteome: history, character, and diagnostic prospects: History, character, and diagnostic prospects. Mol. Cell. Proteomics 1, 845–867 (2002).

5. Uhlén, M. et al. The human secretome. Sci. Signal. 12, 1–9 (2019).

6. Álvez, M. B. et al. A human pan-disease blood atlas of the circulating proteome. Science eadx2678 (2025).

7. Schwenk, J. M. et al. The Human Plasma Proteome Draft of 2017: Building on the Human Plasma PeptideAtlas from Mass Spectrometry and Complementary Assays. J. Proteome Res. 16, 4299–4310 (2017).

8. Geyer, P. E. et al. Plasma Proteome Profiling to Assess Human Health and Disease. Cell Systems 2, 185–195 (2016).

9. Tognetti, M. et al. Biomarker Candidates for Tumors Identified from Deep-Profiled Plasma Stem Predominantly from the Low Abundant Area. J. Proteome Res. 10.1021/acs.jproteome.2c00122 (2022) doi:10.1021/acs.jproteome.2c00122.

10. Aebersold, R. & Mann, M. Mass spectrometry-based proteomics. Nature 422, 198–207 (2003).

11. Nilsson, T. et al. Mass spectrometry in high-throughput proteomics: ready for the big time. Nat. Methods 7, 681–685 (2010).

12. Geyer, P. E. et al. The circulating Proteome─Technological developments, current challenges, and future trends. J. Proteome Res. 10.1021/acs.jproteome.4c00586 (2024) doi:10.1021/acs.jproteome.4c00586.

13. Lou, R. & Shui, W. Acquisition and analysis of DIA-based proteomic data: A comprehensive survey in 2023. Mol. Cell. Proteomics 23, 100712 (2024).

14. Bruderer, R. et al. Analysis of 1508 plasma samples by capillary-flow data-independent acquisition profiles proteomics of weight loss and maintenance. Mol. Cell. Proteomics 18, 1242–1254 (2019).

15. Messner, C. B. et al. Ultra-High-Throughput Clinical Proteomics Reveals Classifiers of COVID-19 Infection. Cell Systems 11, 11–24.e4 (2020).

16. Kamalian, A. et al. Unveiling proteomic and peptide-level modifications in cerebrospinal fluid and plasma in normal cognitive aging. Commun. Med. (Lond.) 5, 474 (2025).

17. Kirsher, D. Y. et al. Current landscape of plasma proteomics from technical innovations to biological insights and biomarker discovery. Commun. Chem. 8, 279 (2025).

18. Beimers, W. F. et al. Technical evaluation of plasma proteomics technologies. J. Proteome Res. 24, 3074–3087 (2025).

19. Al Siblani, S., Armengaud, J. & Lozano, C. Systematic evaluation of depletion and enrichment technologies for platelet-free plasma proteomics. J. Proteome Res. 10.1021/acs.jproteome.5c01056 (2026) doi:10.1021/acs.jproteome.5c01056.

20. Boschetti, E. & Giorgio Righetti, P. Hexapeptide combinatorial ligand libraries: the march for the detection of the low-abundance proteome continues. Biotechniques 44, 663–665 (2008).

21. Blume, J. E. et al. Rapid, deep and precise profiling of the plasma proteome with multi-nanoparticle protein corona. Nat. Commun. 11, 1–14 (2020).

22. Wu, C. C. et al. Enrichment of extracellular vesicles using Mag-Net for the analysis of the plasma proteome. Nat. Commun. 16, 1–15 (2025).

23. Völlmy, F. et al. Comparison of common plasma proteomics workflows reveals distinct preanalytical biases by sample preparation and nanoparticle enrichment. J. Proteome Res. 10.1021/acs.jproteome.6c00048 (2026) doi:10.1021/acs.jproteome.6c00048.

24. Wu, H., Zhang, Y., Wang, Y. & Ouyang, X. Proteonano^TM^: A robust platform for large cohort deep plasma proteomic studies. bioRxiv 2024.08.20.608582 (2024) doi:10.1101/2024.08.20.608582.

25. Bandow, J. E. Comparison of protein enrichment strategies for proteome analysis of plasma. Proteomics 10, 1416–1425 (2010).

26. Palstrøm, N. B., Rasmussen, L. M. & Beck, H. C. Affinity capture enrichment versus affinity depletion: A comparison of strategies for increasing coverage of low-abundant human plasma proteins. Int. J. Mol. Sci. 21, 1–11 (2020).

27. Kverneland, A. H. et al. Benchmarking plasma proteomics workflows and their correlation to clinical routine protein assays. J. Proteome Res. 25, 3188–3200 (2026).

28. Cedervall, T. et al. Understanding the nanoparticle-protein corona using methods to quantify exchange rates and affinities of proteins for nanoparticles. Proc. Natl. Acad. Sci. U. S. A. 104, 2050–2055 (2007).

29. Lundqvist, M. et al. Nanoparticle size and surface properties determine the protein corona with possible implications for biological impacts. Proc. Natl. Acad. Sci. U. S. A. 105, 14265–14270 (2008).

30. Huang, T. et al. Multiplexed nanoparticle protein corona enables accurate and precise deep plasma proteomics. J. Proteome Res. 24, 5884–5893 (2025).

31. Saei, A. A. et al. Leveraging nanoparticle protein corona to advance plasma proteome profiling. Nat. Commun. 17, 5291 (2026).

32. Huang, T. et al. Protein coronas on functionalized nanoparticles enable quantitative and precise large-scale deep plasma proteomics. bioRxivorg 2023.08.28.555225 (2023) doi:10.1101/2023.08.28.555225.

33. Gao, H. et al. Systematic evaluation of blood contamination in nanoparticle-based plasma proteomics. EMBO Mol. Med. 1–22 (2025).

34. Korff, K. et al. Pre-analytical drivers of bias in bead-enriched plasma proteomics. EMBO Mol. Med. 17, 3174–3196 (2025).

35. Geyer, P. E. et al. Plasma Proteome Profiling to detect and avoid sample-related biases in biomarker studies. EMBO Mol. Med. 11, 1–12 (2019).

36. Gao, H., Korff, K., Mann, M. & Guo, T. Deeper is not always better in plasma proteomics. Nat. Biotechnol. 10.1038/s41587-026-03106-3 (2026) doi:10.1038/s41587-026-03106-3.

37. Gronauer, T. F. et al. Blood matrices and sample preparation influence blood marker discovery. J. Proteome Res. 25, 405–417 (2026).

38. Sun, W., Wu, S., Wang, X., Zheng, D. & Gao, Y. A systematical analysis of tryptic peptide identification with reverse phase liquid chromatography and electrospray ion trap mass spectrometry. Genomics Proteomics Bioinformatics 2, 174–183 (2004).

39. Muntel, J. et al. Biomarkers. Alzheimers. Dement. 21 Suppl 2, e100311 (2025).

40. Heeb, L. et al. Abstract 106: Integrated plasma multi-omics profiling identifies circulating predictive biomarkers and biological pathways associated with treatment response in NSCLC. Cancer Res. 86, 106–106 (2026).

41. Rigutto, A. et al. Abstract 3306: Comprehensive plasma profiling in NSCLC patients undergoing IO- and chemotherapy identifies potential proteomic signatures for progression-free survival. Cancer Res. 85, 3306–3306 (2025).

42. Rigutto, A. et al. 242P: SAKK 17/18 ORIGIN trial: Efficacy and safety from a multicentre phase II trial of gemcitabine and atezolizumab in pretreated patients with pleural mesothelioma. J. Thorac. Oncol. 20, S152–S153 (2025).

43. Järvinen, E., Liu, X., Varjosalo, M. & Keskitalo, S. Automated Mag-Net enrichment unlocks deep and cost-effective LC-MS plasma proteomics. J. Proteome Res. 25, 618–632 (2026).

44. Batth, T. S. et al. Protein Aggregation Capture on Microparticles Enables Multipurpose Proteomics Sample Preparation. Mol. Cell. Proteomics 18, 1027–1035 (2019).

45. Bruderer, R. et al. Extending the limits of quantitative proteome profiling with data-independent acquisition and application to acetaminophen-treated three-dimensional liver microtissues. Mol. Cell. Proteomics 14, 1400–1410 (2015).

46. Cox, J., Hein, M. Y., Luber, C. a. & Paron, I. Accurate proteome-wide label-free quantification by delayed normalization and maximal peptide ratio extraction, termed MaxLFQ. Mol. Cell. Proteomics 13, 2513–2526 (2014).

47. Huang, T. et al. Combining precursor and fragment information for improved detection of differential abundance in data independent acquisition. Mol. Cell. Proteomics 19, 421–430 (2020).

48. Storey, J. D. A direct approach to false discovery rates. J Royal Stat Soc Ser B-Stat Method 64, 479–498 (2002).

49. Pino, L. K. et al. Matrix-matched calibration curves for assessing analytical figures of merit in quantitative proteomics. J. Proteome Res. 19, 1147–1153 (2020).

50. Wahle, M. et al. A novel hybrid high-speed mass spectrometer allows rapid translation from biomarker candidates to targeted clinical tests using 15N-labeled proteins. Mol. Cell. Proteomics 24, 101050 (2025).

51. Jornitz, M. W. & Meltzer, T. H. Sterile Filtration: A Practical Approach. (CRC Press, Boca Raton, FL, 2000).

52. Genovese, B. N. et al. Specimen inactivation methods for Proteomics─Comparisons of irradiation, chemical, and heat treatments on downstream serum analyses. J. Proteome Res. 24, 5282–5291 (2025).

53. WHO Guidelines on viral inactivation and removal procedures intended to assure the viral safety of human blood plasma products, Annex 4, TRS No 924. https://www.who.int/publications/m/item/WHO-TRS924-Annex4.

54. Keerthikumar, S. et al. ExoCarta: A Web-based compendium of exosomal cargo. J. Mol. Biol. 428, 688–692 (2016).

55. Salem, I. et al. Methods for extracellular vesicle isolation from cancer cells. Canc. Drug Resist. 3, 371–384 (2020).

56. Krusen, B. M. et al. Improving a plasma biomarker panel for early detection of pancreatic ductal adenocarcinoma with aminopeptidase N (ANPEP) and polymeric immunoglobulin receptor (PIGR). Clin. Cancer Res. 32, 756–769 (2026).

57. Wang, Z. et al. The prognostic and clinical value of CD44 in colorectal cancer: A meta-analysis. Front. Oncol. 9, 309 (2019).

58. Baig, N. et al. Clinical significance and comparison of flotillin 1 expression in left and right colon cancer. Oncol. Lett. 18, 997–1004 (2019).

59. Li, T., Cao, C., Xiong, Q. & Liu, D. FLOT2 overexpression is associated with the progression and prognosis of human colorectal cancer. Oncol. Lett. 17, 2802–2808 (2019).

60. Li, Z., et al. Periostin expression and its prognostic value for colorectal cancer. Int. J. Mol. Sci. 16, 12108–12118 (2015).

61. Gargalionis, A. N., Papavassiliou, K. A. & Papavassiliou, A. G. Targeting STAT3 signaling pathway in colorectal cancer. Biomedicines 9, 1016 (2021).

62. Deutsch, E. W. et al. The State of the Human Proteome in 2014/2015 as viewed through PeptideAtlas: enhancing accuracy and coverage through the AtlasProphet. J. Proteome Res. 150703045314008 (2015).

